# Force Scaling in Active Swarm of Microtubules via Magnetic Manipulation

**DOI:** 10.1101/2025.11.20.689561

**Authors:** Mst. Rubaya Rashid, Mousumi Akter, Arif Md. Rashedul Kabir, Kazuki Sada, Tetsuya Hiraiwa, Akinori Kuzuya, Ibuki Kawamata, Marie Tani, Akira Kakugo

**Author notes:** Akira Kakugo **Email:**. **Author Contributions:** Conceptualization: A. Kakugo; methodology: M.R.R., M.A.; investigation: M.R.R.; visualization: M.R.R; funding acquisition: A. Kakugo, A.M.R.K., and T.H.; project administration: M.R.R, M.A., A.M.R.K., K.S., and A. Kakugo; supervision: A. Kakugo; writing: M.R.R; writing—review and editing: M.R.R., M.A., A.M.R.K., A. Kuzuya, T.H., K.S., I.K., M.T., and A. Kakugo. **Competing Interest Statement:** The authors declare that they have no competing interests. **Classification:** Physical Sciences- Biophysics and Computational Biology.

## Abstract

Scalability refers to the ability of a system to enhance performance with increasing size. This ability is a defining feature of natural swarms such as ant and bee colonies. Extending this principle to artificial active matter has motivated the creation of molecular swarms composed of myosin-driven actin filaments and kinesin-driven microtubules (MTs). While these swarms exhibit cooperative transport behaviors that surpass the abilities of individual filaments, their collective force generation has remained unquantified. Here, we introduce a straightforward electromagnetic tweezer approach to directly measure the forces generated by MT swarms propelled by surface-bound kinesin motors. We find that force output changes with swarm size, demonstrating quantitative scalability and linking collective organization to mechanical performance. These results establish MT swarms as a model system for scalable, force-producing active matter and provide a foundation for designing biomolecular devices that exploit collective dynamics for microscale actuation and transport.

**Significance Statement:** The cooperative activity of multiple motile agents is observed in nature, known as swarming. In artificial systems, swarming has been demonstrated using active agents like kinesin and MTs, successfully enabling efficient cargo transportation. However, the force generated by the swarm remains unexplored, which is one of the keys to understanding the principles underlying swarm systems. This study introduces a simple method employing electromagnetic tweezers and magnetic beads to measure the force of the MT swarm accurately. The measured forces within the microtubule swarms of different sizes unveil the scaling effect, the additive nature of force generation as the kinesin unit increases. The force study sheds light on the fundamental properties, work performance, efficiency, and execution of the MT swarm.

## Introduction

Swarming is a striking natural phenomenon in which the collective behavior of motile entities enables tasks far beyond the capacity of individuals ^1^. Ant and bee colonies exemplify how scalability, robustness, and efficiency emerge from simple agents acting together ^2–7^. Inspired by these natural advantages, researchers have engineered artificial swarms at scales ranging from macro to micro to harness collective performance for technological applications ^8–11^.

Recent technological progress has enabled the miniaturization of machines, amplifying their efficacy in diverse nanotechnological applications ^12,13^. Among these microscale machines, chemically fueled biomolecular motor systems, such as microtubule (MT)– kinesin/dynein and actin–myosin, have garnered significant interest due to their distinctive motion characteristics and engineering potential ^14–21^. In particular, swarms of kinesin-propelled MTs have been extensively studied ^22–26^. Advances in DNA nanotechnology have fine-tuned MT swarm behavior through the manipulation of intermolecular interactions, providing a versatile interface for programmable control ^16,22,27^. DNA enables logic operations, systematic optimization, photo-responsiveness, and size-tunable assemblies, positioning MT swarms as programmable molecular devices and robotic systems^22,28,29^.

Importantly, MT–kinesin swarms exhibit superior cooperative transport compared to individual filaments, excelling in the quantity and size of cargo they can carry as well as the distances they traverse ^28^. The collective forces produced by hundreds of kinesin-propelled MTs may parallel the coordinated force sharing observed in natural swarms, such as the cooperative transport of ants ^4,5,7^, where group performance is tightly linked to scalable force output. Efficient force generation and its scalability are therefore pivotal metrics for future applications.

Single kinesins generate forces of about 6–8 pN and have been extensively studied. Force generated by single kinesins or small ensembles up to ten motors have also been measured using techniques such as optical tweezers ^30–36^, atomic force microscopy ^37^, DNA linkers ^38^, contractile active networks ^39^, and magnetic tweezers ^40–44^. However, the integrated force generation of MT swarms remains unquantified, as measuring the collective output of hundreds of motors is hindered due to the difficulty of controlling many MTs at once. Electromagnetic tweezers, with their broad tunable force range ^40–44^, could provide a promising approach to overcome this limitation and enable quantitative investigation of scalable force generation within MT swarms.

We quantified the collective forces generated by DNA-tethered microtubule (MT) swarms propelled by thousands of kinesins using a customized electromagnetic-tweezer system integrated with magnetic beads. By constraining the swarms into ring geometries with uniform polarity rather than bundles ^22^, we achieved stable and reproducible force measurements. The collective output scaled linearly with swarm size, producing forces nearly two orders of magnitude greater than those generated by a single kinesin (6–8 pN) ^30–34^. This scalable and tunable force generation demonstrates MT swarms as a robust model of collective motor activity and a promising platform for biomolecular devices and synthetic active-matter applications.

## Results

### Concept

The MT swarms in a ring configuration circulate at a constant velocity, powered by hundreds of kinesin motors converting the chemical energy of ATP into mechanical work (Figure 1A). While each kinesin acts individually, their collective activity generates the integrated force of the swarm. To directly quantify this swarm force, *F*_*sw*_, we developed a custom electromagnetic tweezer (EMTw) system coupled with fluorescence microscopy.

**Figure 1.**
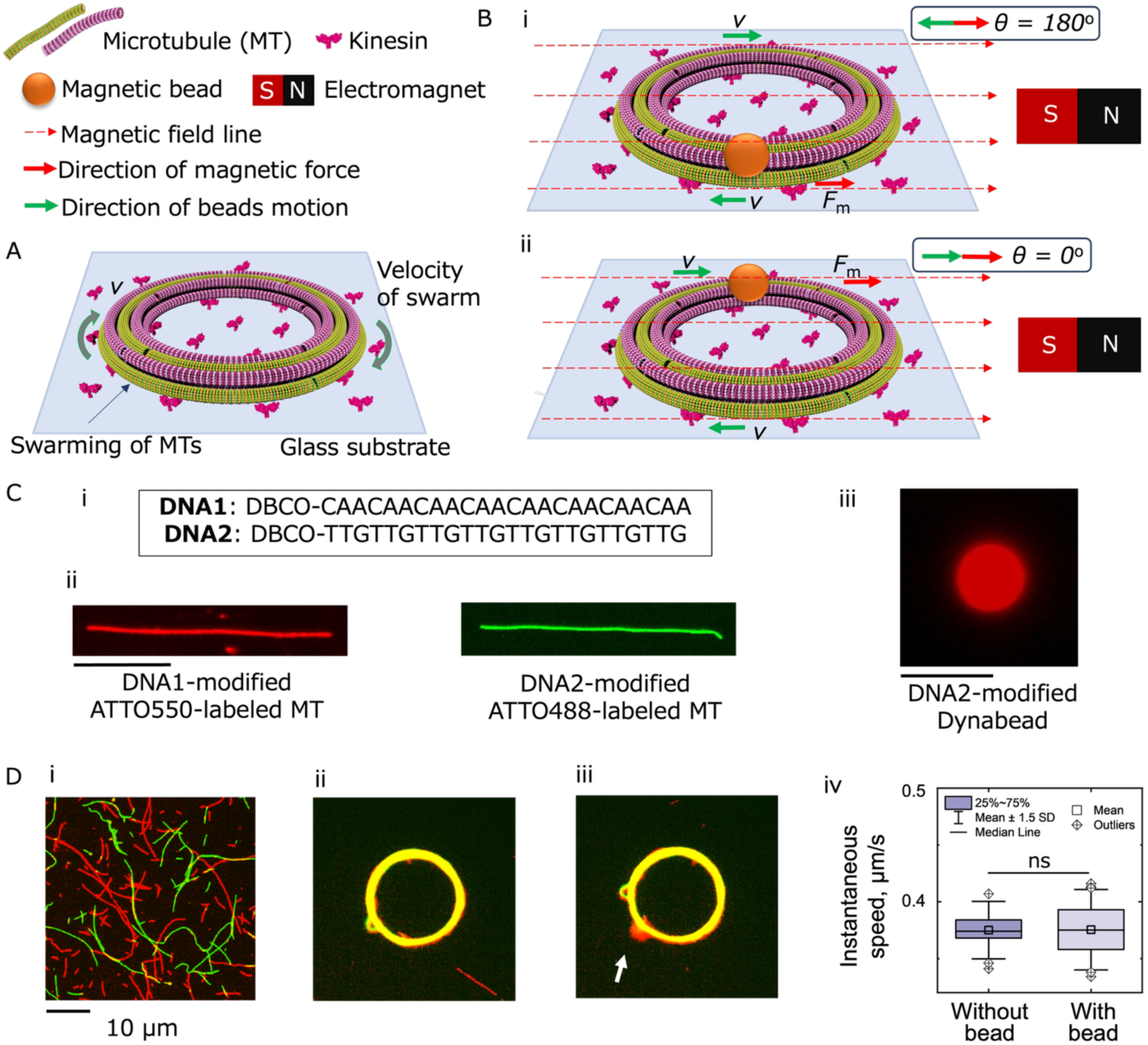
Concept, preparation of swarm rings, and bead loading; (A) Schematic illustration of force generation by microtubule (MT) swarms propelled by multiple kinesins (n >> 10), where n denotes the number of kinesin motors. (B) Experimental setup for determining the force of a circular MT swarm using a magnetic bead and an EMTw: (i, ii) schematic representation of the model. θ denotes the angle between the direction of the applied magnetic force and the direction of bead motion. (C) DNA design and swarm formation: (i) DNA sequences used to assemble MT swarms and attach magnetic beads, fluorescence microscopic images of (ii) MT swarm units, scale: 10 μm, and (iii) a bead, scale: 5 μm. (D) Fluorescence microscopy images of kinesin-propelled MTs: (i) single MT state, (ii) swarming state, and (iii) swarm attached to a bead, scale bars = 10 μm. (iv) Box plot of instantaneous MT swarm velocities with and without bead attachment. A total of n = 150 instantaneous velocity measurements were analyzed across both conditions. Outliers and mean ± 1.5 SD are indicated. Paired-sample t-test shows no significant difference between the two datasets (p > 0.01; ns).

We chose ring-shaped swarms over bundle-shaped swarms to ensure reliable force measurements. Bundle swarms translate randomly and frequently shift position ^22,23^, complicating force calibration in the nonuniform EMTw field. By contrast, ring swarms circulate around a fixed center of mass, providing a stable trajectory that allows repeated and reproducible force measurements ^45–48^.

In our setup, Figures 1A and B, a magnetic bead tethered to the MT ring via DNA hybridization, serves as a force probe circulating with the swarm. The collective force generated by multiple kinesins, *F*_*sw*_, (green arrow in Figure 1A) propels the MT swarm, while the applied electromagnetic force (EMF) adds an additional magnetic force, *F*_*m*_, (red dashed arrow in Figure 1B) acting on the bead-loaded ring.

The bead’s direction of motion changes continuously along the ring, as illustrated in Figure 1B(i, ii), where it alternates between opposing and aligning with the direction of applied EMF. Given this changing orientation, the effective net force on the bead, defined as the tangential magnetic force, *F*_m_^θ^., effectively opposes or assists the swarm. As the bead circulates, the tangential magnetic force, *F*_m_^θ^ varies continuously, leading to periodic modulations in swarm velocity. These oscillations arise from the interplay between the kinesin-generated force *F*_sw_ and the applied magnetic force *F*_m_. In principle, by calibrating the relative contributions of these two forces, the collective force of the MT swarm can be determined. Under our conditions, viscous drag was negligible, as discussed later.

### Construction of the swarm system

The MT swarms were constructed through sequential MT polymerization, DNA modification, and assembly of MTs on kinesin-coated surfaces (Figure 1C–D). First, MTs were polymerized from tubulin mixtures containing 80% azide-functionalized subunits (for subsequent click reaction with DBCO-DNA) and 20% fluorophore-labeled tubulins (ATTO550 and ATTO488, used to generate MTs of two distinct colors for fluorescence imaging) in the presence of guanosine triphosphate (GTP) (Figure S1A) ^49,50^.

Two complementary DNA strands, DNA1 and DNA2 (Table S1), were conjugated to the MTs via azide–alkyne cycloaddition reaction, resulting in two distinct sets of DNA-modified MTs (Figure 1C(i–ii)) ^49,50^. Superparamagnetic Dynabeads (beads) were subsequently functionalized with DNA2 (Figure 1C(iii), Figure S1B), enabling their hybridization-based tethering to DNA1-modified MTs and incorporation into the MT swarms.

For swarm assembly, a flow cell was coated with his-tagged 400 nM human kinesin-1 motors (recombinant kinesin, consisting of the first 573 amino acid residues of human kinesin 1) ^51^. DNA1- and DNA2-modified MTs were then infused into the kinesin-coated flow cell, where they began gliding along the kinesin surface powered by ATP hydrolysis (Figure 1D(i)) ^49,50^. Adjacent MTs of the same polarity associated through hybridization of the complementary 24-base DNA1 and DNA2 strands, while the intrinsic flexibility of MTs promoted their spooling into stable ring-shaped swarms (Figure 1D(ii)) ^52^. These rings circulated clockwise or counterclockwise.

After 30–40 minutes, when swarm formation reached about 60–70% (Figure S2), DNA2-functionalized beads dispersed in ATP buffer were introduced into the flow cell, where they tethered to the swarm by DNA hybridization (Figure 1D(iii), Figures S3–S4). To prevent nonspecific interactions, free-floating beads were thoroughly washed out of the chamber, ensuring specific attachment.

The circulating velocity of the swarm rings remained unchanged upon bead attachment (0.38 ± 0.02 μm/s vs. 0.38 ± 0.03 μm/s, mean ± SD, n = 25; Figure 1D(iv)), indicating that the bead load did not perturb swarm dynamics. Minor variations across experiments were observed (Fig. S5, S9 (inset)), which may reflect temperature fluctuations, timing effects following ATP addition, and other uncontrolled variables.

### Establishment and Calibration of a Custom EMTw

To directly apply calibrated forces to MT swarms, we developed a custom EMTw integrated with a fluorescence microscope (Figure 2A, Fig. S6). Prior to swarm force measurements, calibration was performed using superparamagnetic beads suspended in 80% glycerol (v/v; viscosity ∼0.073 Pa·s at room temperature), which reduced bead velocity and counteracted gravitational settling.

**Figure 2.**
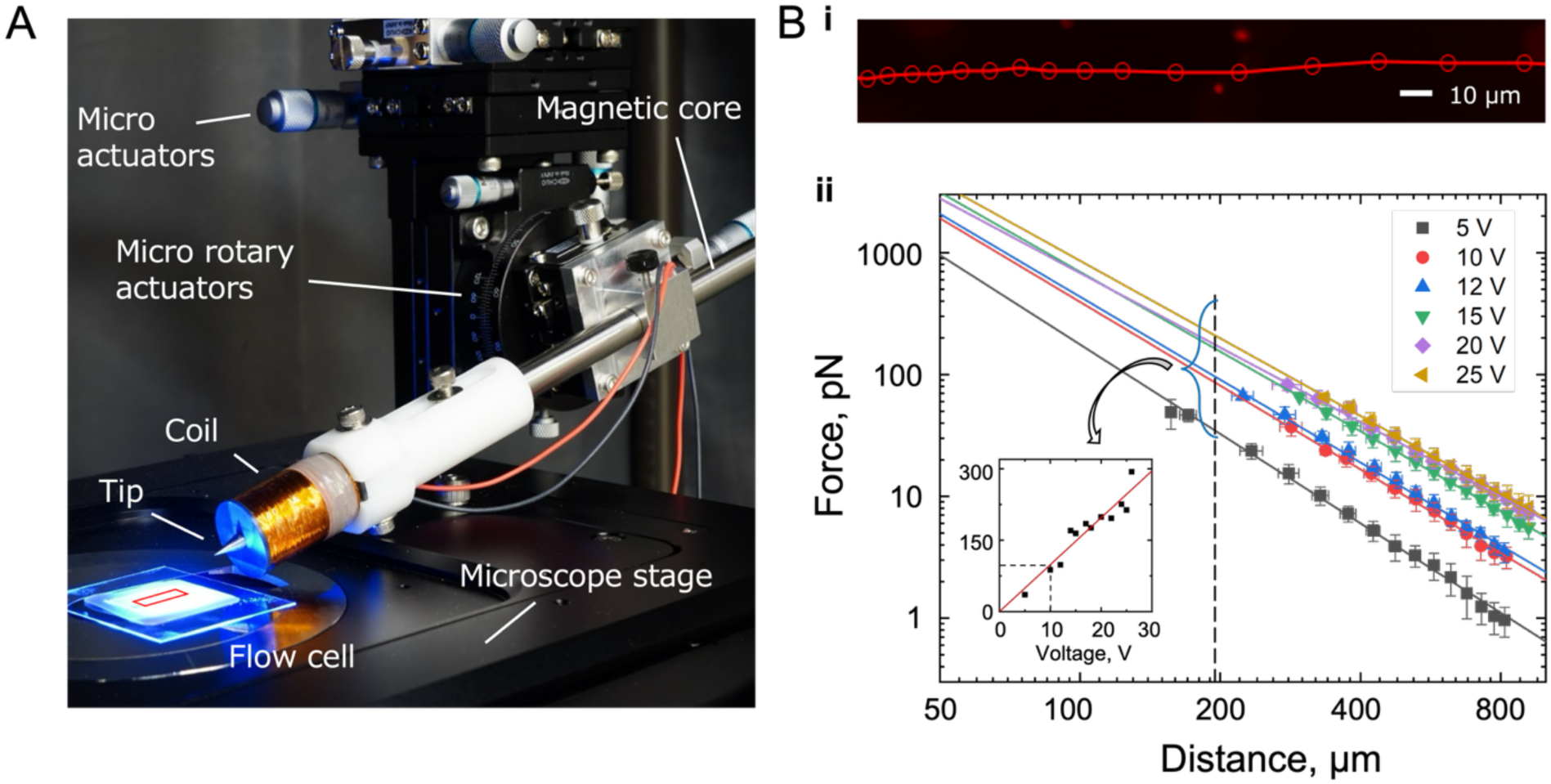
Force calibration of Dynabeads using the custom-built electromagnetic tweezer (EMTw) integrated with a fluorescence microscope. (A) Photograph of the EMTw setup and different parts. (B.i) Representative bead trajectory under applied EMF. (B.ii) Log–log plot of force versus distance at different voltages; error bars represent standard deviations. Data were fitted with a power law (eq. ii). Inset: linear dependence of force on voltage at a fixed distance, enabling precise force calibration at given voltage–distance conditions.

Beads dispersed in glycerol were introduced into the flow cell, and their time-lapse trajectories under the applied EMF were recorded using fluorescence microscopy (Figure 2B(i)). As shown in Figure 2B(i), the beads translated laterally while remaining in focus, confirming gravitational and magnetic forces in the vertical direction were balanced. In our experimental setup, the Reynolds number was significantly low, meaning that viscous friction of the solvent dominated over inertial effects. The viscous drag force on the beads, *F*, was therefore estimated using Stokes’ law as:

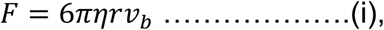

where *η* is the medium viscosity, *r* is bead radius, and 𝑣_*b*_ is the velocity determined from the trajectory of the free beads applying EMF ^53^.

Force–distance (F–d) curves for beads at varying voltages shows that forces increases as beads approach towards the EMTw’ tip, consistent with the stronger magnetic field gradient at shorter distances (Figure 2B(ii)). Force and velocity were optimized by adjusting applied voltage and bead-to-EMTw distance; error bars show standard deviations.

The straight lines in Figure 2B(ii) were obtained by fitting the experimental data for each applied voltage V with a power-law function with,

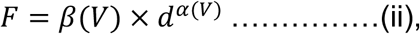

which is a rearrangement of the previously established relation *F* = *F*_0_(*d*/*d*_0_)^*c*(*I*)^ ^53^ with *α*(*V*) = *c*(*I*) and 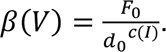 As *F*_0_ and *d*_0_ cannot be directly determined in our system, the derivation is provided in the SI. As shown in the inset of Figure 2B(ii), the extracted forces at a fixed distance (vertical dashed line) scaled linearly with applied voltage (*F* ∝ *V*), enabling straightforward calibration of the EMTw system. This linear fitting provides a reliable means to determine force at given voltage–distance conditions, ensuring precise and tunable force control.

### EMF Application to Bead-Attached Swarms and Force Determination

Fluorescence microscopy confirmed the attachment of a single bead to the MT swarm ring, exhibiting circular velocity. The EMTw tip was positioned about 100–300 μm from the swarm, and EMF was applied as an external force to modulate its circular rotation. As the applied EMF increased to values comparable with the swarm force, the swarm velocity was modulated in a manner dependent on the bead’s position and direction of motion (Figure 3, Movie 1).

**Figure 3.**
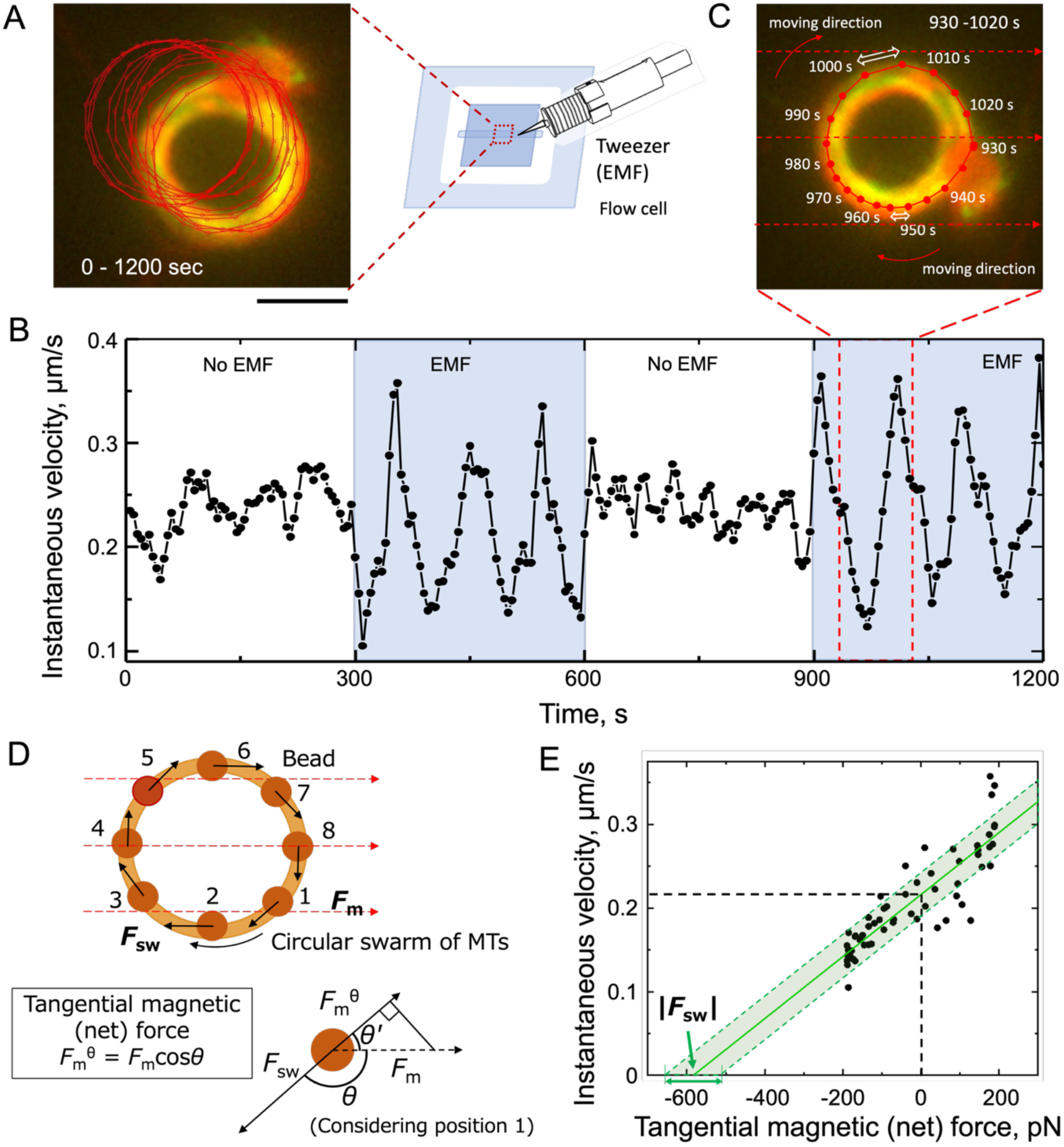
Controlling the velocity of a bead-attached circular swarm by electromagnetic tweezer (EMTw) and force estimation from velocity scaling. (A) Trajectory of the bead-attached swarm ring without EMF (0 V, 0–300 s and 600– 900 s) and with EMF (10 V, 300–600 s and 900–1200 s). Scale bar: 5 µm. (B) Instantaneous velocity of the swarm ring without EMF (average velocity, no shade) and with EMF (periodic velocity changes, blue shade). (C) Single-cycle trajectory (930–1020 s) of the bead under EMF. Displacement per interval varied depending on the relative orientation between the applied magnetic field and bead motion. (D) Schematic of bead positions during circulation. Top: the attached bead changes its direction of motion, *θ* with the circulating swarm, while the magnetic field lines remain fixed. Bottom: Tangential magnetic force (net force), 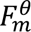 on the bead at each position, determined by the angle, *θ* between *F*_*m*_ and *F*_*sw*_. (E) The velocity |𝑣| as a function of the tangential magnetic force 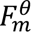 on the bead. Black dashed line: average velocity of the swarm ring at zero applied force. Green solid line: fit to eq. (v) (*R*^2^ ≈ 0.79). Extrapolation to zero velocity yields the swarm force |*F*_*sw*_|, (green single-headed arrow). Error bands are shown as a green shadow, with standard deviation indicated by the double-headed arrow.

In a representative case (Figure 3, Movie 1), the swarm ring had an outer diameter of ∼9 μm and a width of ∼1.4 μm, measured from fluorescence images. The bead-attached ring rotated at |𝑣| = 0.27 ± 0.02 μm/s in the absence of EMF. The distance between the EMTw tip and swarm center, estimated from full-frame fluorescence images (276 μm × 232 μm), was ∼177 μm, far larger than the 9 μm ring diameter. Under this condition, the magnetic field across the swarm can be approximated as parallel with only ±1° angular deviation (Figure S7). For simplified analysis, the tip-to-swarm distance was assumed constant, despite the bead circulating within the annulus. Vertical force balance was maintained under the same conditions as in the calibration experiment.

The swarm motion was monitored for 20 minutes, alternating between time segments with and without EMF: 0–300 s (no EMF), 300–600 s (with EMF), 600–900 s (no EMF), and 900–1200 s (with EMF) (Figure 3A). The instantaneous velocity of the bead-attached swarm was extracted from fluorescence movies. In the absence of EMF, the velocity remained relatively stable, whereas under EMF it varied periodically (Figure 3B, Fig. S8). Figure 3C shows a representative one-cycle trajectory of a bead under EMF (930–1020 s), with the corresponding schematic in Figure 3D, where red dashed lines indicate the electromagnetic field lines.

The external magnetic force, *F*_*m*_, applied through the bead was determined from the force calibration curve and corresponding bead position (Figure 2B). The net force, defined as the tangential component of the magnetic force acting on the bead, varies as the bead’s direction of motion changes with the circulating swarm ring (Figure 3D), whereas the direction and magnitude of the applied electromagnetic field remain constant. The net force is estimated as:

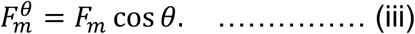

Here, *θ* is the angle between the bead’s direction of motion and the magnetic field lines. Depending on *θ* and the bead’s position on the swarm ring, the net forces 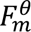 can either assist (positive) or oppose (negative) the swarm force *F*_*sw*_, which remains approximately constant; as a result, the bead’s velocity varies accordingly.

The smallest and largest displacements in one rotation cycle were observed at 950 – 955 s and 1000 – 1005 s, respectively (Figure 3C). At 950–955 s (*θ* ≅ *π*, field opposing motion), the displacement was smaller than average, whereas at 1000 – 1005 s (*θ* ≅ 0, field aligned with motion), the displacement was larger. As shown in Figure 3E, the instantaneous velocity |𝑣| increased with the tangential magnetic force, 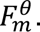

We describe the motion of the bead attached to the MT swarm under low Reynolds number conditions, where inertia is negligible. Consequently, the total force on the bead, 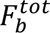 can be approximated as zero:

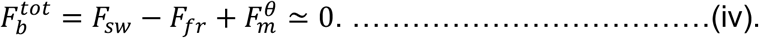

Here, 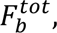 is the sum of the driving force generated by the MT swarm *F*_*sw*_, the viscous frictional force *F*_*fr*_, and the effective tangential magnetic force 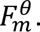 The viscous drag on the bead, (*F*_*η*,*b*_ = 6 *πηr*𝑣_*b*_), is on the order of 10 fN and was neglected because it is much smaller than the applied magnetic force 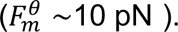 Notably, this viscous contribution is smaller than in the calibration experiment performed in a glycerol medium (Figure 2B).

We estimated the driving force *F*_*sw*_ and frictional force *F*_*fr*_ from the force generated by a single kinesin *f*_*kin*_ and the corresponding frictional force *f*_*fr*_, such that: *F*_*sw*_ = *f*_*kin*_*N*_*kin*_ and *F*_*fr*_ = *f*_*fr*_*N*_*kin*_. Here *N*_*kin*_ is the number of involved kinesins. Since the velocity of a kinesin on a MT, 𝑣_*kin*_ decreases linearly with the applied load ^30,54^, it is natural to assume that kinesin experiences a frictional force proportional to its velocity. Here, we assume that the kinesin-driven swarm velocity 𝑣_*kin*_ is comparable to the bead’s velocity 𝑣_*b*_, i.e. 𝑣_*kin*_ ≈ 𝑣_*b*_, because the bead is attached to the swarm propelled by kinesins. Therefore, the frictional force can be expressed as *f*_fr_ = *k*_𝑣_𝑣_*b*_, where the prefactor *k*_𝑣_ is on the order of 10^−;30^. Accordingly, the total frictional force exerted on the bead by the MT swarm can be expressed as, *F*_*fr*_ = (*k*_𝑣_𝑣_*b*_) *N*_*kin*_. Substituting this into eq. (iv), the equation can be rewritten as:

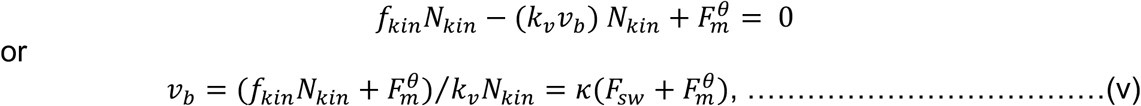

where *κ* = 1/*k*_𝑣_*N*_*kin*_, whose order is about 10^2^ m/Ns.

The experimental data (Figure 3E) are well described by the linear function in Eq. (v), yielding a fitting parameter of *κ* ≈ 3.7 × 10^2^ m/Ns (solid line). From the x-intercept of the extrapolated fit, corresponding to the force required to stop the swarm, we obtained the swarm force as *F*_*sw*_ = 583 ± 80 pN, where the uncertainty represents the standard error.

### Force generation across swarm rings of varying size

Forces of swarm rings with different diameters and widths were estimated from multiple events. The measured forces ranged from a few hundred to several thousand piconewtons (pN), depending on swarm size and geometry (length, width, and height). Because wider and larger rings occupy larger areas of the kinesin-coated surface, they involve more motors and thus yield higher forces. Additional representative events are provided in Fig. S9.

To further analyze force behavior, we quantified the average width and diameter of swarm rings using thresholded fluorescence images (Fig. S10). The results indicate that the swarm force (*F*_*sw*_) generally increases with the width and diameter of MT rings, although variability becomes larger as width and diameter increase. To capture this trend more directly, we calculated the annulus area of each swarm ring (from diameter and width, excluding the inner void) and plotted the force against the annulus area (Figure 4A). The force shows an overall increasing trend with annulus area, consistent with the involvement of more MTs and kinesins in larger swarm rings.

**Figure 4.**
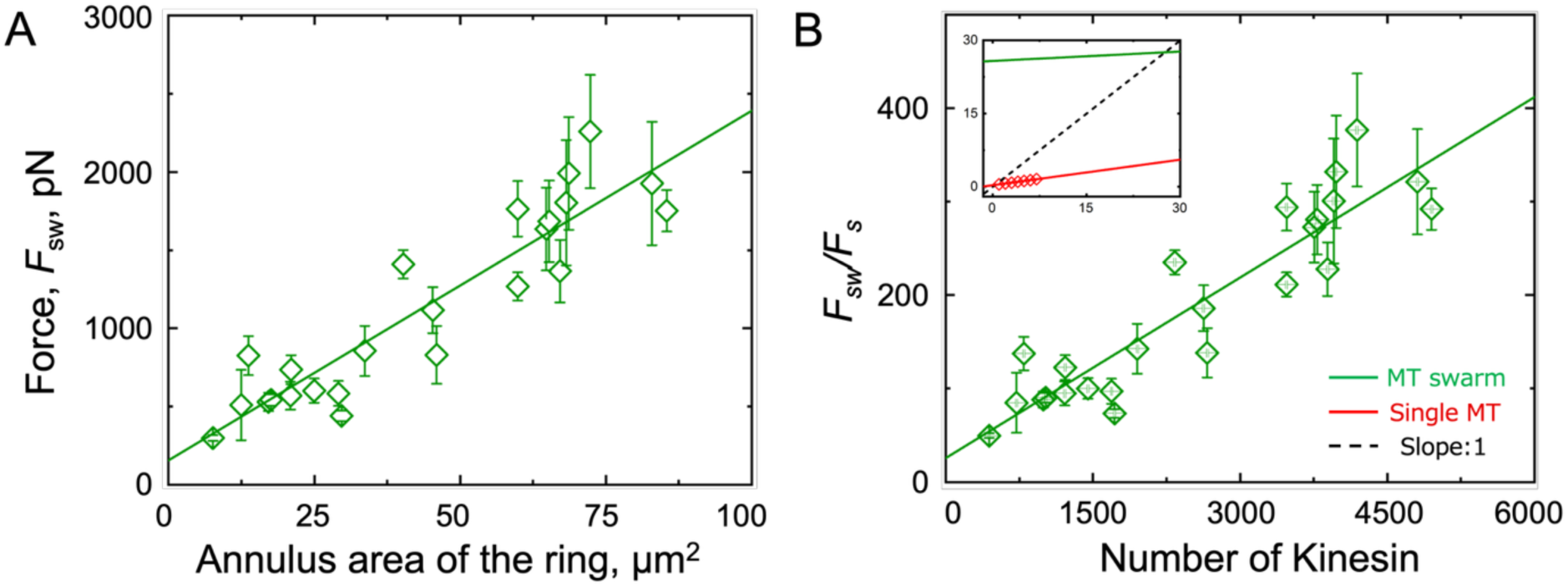
Force generation by MT swarms. (A) Relationship between swarm force, *F*_*sw*_, and annulus area of individual MT swarm rings. The ring width was determined from fluorescence images converted to threshold binary images. *F*_*sw*_, increases with the occupied swarm area, and error bars represent standard errors from 26 independent measurements. (B) Comparison of swarm forces, *F*_*sw*_, involving a large number of kinesins, with single-MT motility driven by a few kinesins (∼10 motors, inset; Fig. S15). The forces, *F*_*sw*_, (green) and the single MT’s force (red), normalized by the stall force of single kinesin *F*_*s*_, are plotted as a function of the number of kinesins. The dashed black line denotes the theoretical slope of 1, representing purely additive force contributions from individual kinesins. For single-MT motility, (inset, red line, data obtained from ref. ^58^, the rescaled force increases linearly with kinesin number, yielding a slope of 0.17 ± 0.01 (R² = 0.90). For swarm rings, the linear fit produced a slope of 0.07 ± 0.01 (R² = 0.79). Therefore, the fitted curves highlight the linear increase in force generation with kinesin number for both swarm rings (green line) and single-MT motility (inset, red line).

### Estimating MT and kinesin numbers

Larger swarms, with greater annulus areas, involve more MTs and kinesins than smaller ones. To estimate MT numbers, we relied on a previous study ^55^. In that study, photo-dissociation of MT swarms using photo-responsive DNA and UV light, observed via fluorescence microscopy, was validated against height measurements from high-speed AFM (Fig. S11A). Swarm ring widths from the current force experiments were compared with prior fluorescence-based measurements (Fig. S11B), confirming that wider rings contained more MT filaments and contributed to the observed force range. Kinesin surface density was determined from landing-rate experiments ^56^. At 400 nM kinesin concentration, the density was estimated as 61.05 ± 22.04 μm^−2^ (Figs. S12, S13; Methods). By combining the kinesin density (from landing-rate experiments) ^56,57^ with the annulus area of swarm rings obtained from thresholded binary images, we estimated the total number of kinesins engaged in each ring.

The estimated forces as a function of kinesin number along the annulus area of the swarm rings are shown in Fig. S14. Swarms composed of hundreds of microtubules and driven by 600–5000 kinesins generated forces ranging from approximately 250 to 2500 pN, with the total force increasing with both swarm size and kinesin number, reflecting the collective contribution of motors distributed along the annular region. Such high force-generation capacity and its scalability are critical for the effective implementation of MT swarms, enabling efficient, robust, and cooperative behavior across diverse experimental and practical applications.

### Scaling relative to single kinesins

To assess efficiency, swarm forces, ***F***_***sw***_, were normalized by the stall force of a single kinesin motor (***F*_*s*_** ≈ 6 pN: from ref. ^32^). The normalized force (*F*_*sw*_/*F*_*s*_) was plotted against the estimated number of kinesins (Fig. 4B). The data were well described by a linear fit, indicating that forces scaled with the number of engaged motors for both swarms and single MT motility (Fig. 4B, inset; green and red lines, Fig. S15), consistent with the inherent characteristics of the cooperative action of kinesin motors.

For single MT motility, a slope of 0.17 is obtained from ref. ^58^ and is illustrated by the red line in Fig. 4B (lower scale shown in Fig. S15). This slope is smaller than the unit slope corresponding to strict additivity (black dashed line). For swarm rings, the slope decreases to ∼0.07, likely due to the ring geometry; this deviation will be discussed in the later section.

In Figure 4B (Fig. S15), the intercept of the linear fit for the swarm rings (21.67 ± 25.19) is substantially higher than that obtained for single-MT motility involving only a few kinesins (0.33 ± 0.01). Direct comparison of slope values is difficult, as the experimental conditions and number of engaged motors differ between systems. With increasing kinesin numbers, slope values may decline gradually, from single filaments (a few motors) to large swarms (hundreds to thousands), and at very high counts, force generation may plateau or follow a logarithmic trend rather than linear growth. At present, there is no experimental evidence to fully address this issue; it remains an open question, and systematic future experiments will be necessary to clarify how collective kinesin action transitions from single MTs to swarms.

## Discussion

Our study examines force generation by DNA-tethered, kinesin-propelled MT swarms. In these swarms, we measured forces in the range of 250–2500 pN for motor numbers of approximately *N*_kin_≈ 600–5000, whereas single-MT motility produces only 6–15 pN (*N*_kin_ ≈ 2–10). These results indicate that total force output increases with the number of participating MTs and motors. For context, the forces measured here fall within the ranges, and in some conditions exceed those reported for multi-kinesin and bead-bundle assays ^59,60^.

Although force generation scales linearly with kinesin number in both single-filament and swarm systems, the slope is smaller in the ring geometry. This deviation reflects geometric constraints, as kinesins arranged along a circular path contribute vectorially rather than purely additively. Similar geometry-dependent vectorial summation of forces has been reported in various motor–filament systems, including kinesin moving along bent microtubules ^61^, actomyosin networks ^62^, dynein ensembles ^63^, and confined filament motor vortices ^64^. Together, these examples suggest a general physical rule of collective motor behavior and highlight geometry as a key design factor that governs overall performance.

Within the broader context of molecular robotics, our findings complement ongoing efforts to harness biomolecular motors for engineered actuation. Recent advances include the development of multicellular robotic systems that emulate neural networks ^10,65–67^, and the creation of molecular “artificial muscles,” where DNA-modified MTs hierarchically self-assemble into aster-like structures and larger aggregates ^68^. These systems exhibit dynamic, chemically driven contraction, highlighting the potential of biomolecular motors for engineered actuation. Light-responsive MT–kinesin active networks have also demonstrated macroscopic deformation and programmable spatiotemporal control ^39,69^. Building on these developments, the present work provides direct, quantitative force measurements that link nanoscale motor coordination to mesoscale mechanical performance. By demonstrating both reproducibility and scalability, our DNA-mediated MT swarms establish a robust experimental platform for probing cooperative dynamics and energy transduction in active-matter systems.

In this emerging field, molecular swarm systems stand out for their scalability, parallelism, robustness, and flexibility ^1–3,13,19^. Chemically fueled biomolecular motors, particularly MT–kinesin and MT–dynein assemblies, are of special interest for their rich motion characteristics and engineering potential ^15,16,20^. In such systems, DNA and its derivatives often function as sensors and processors, while MT–kinesin serves as the actuator, enabling programmable molecular-scale robots to collaborate in organized ways ^22,27,50^. By directly quantifying force generation, our work indicates that MT swarm forces scale with motor number and are influenced by spatial organization. These results suggest considerations for designing cooperative molecular systems in which motor number and geometry contribute to mechanical performance.

Looking ahead, integrating this swarm-based actuation with feedback control or chemical signaling could enable adaptive micromachines that sense and respond to their environment. Coupling EMTw-based force measurement with real-time fluorescence or FRET reporters would further link mechanical output to biochemical state, enabling mechanochemical mapping at the single-swarm level. On the applied side, translating these scalable force-generation principles into DNA-origami scaffolds, surface-patterned microdevices, or synthetic organelles could enable advances in targeted drug delivery, nanoscale cargo transport, and responsive soft robotics. In summary, the quantitative relationships, which we identify among (1) motor number, (2) geometry, and (3) collective force output, clarify the advantage of MT swarms as a useful model for cooperative motor mechanics and as a potential platform for future biomolecular actuation strategies.

## Materials and Methods

### Purification of Tubulin and Kinesin

Tubulin was purified from porcine brain using high-concentration PIPES buffer (1 M PIPES, 20 mM EGTA, 10 mM MgCl₂) and stored in BRB80 buffer (80 mM PIPES, 1 mM EGTA, 2 mM MgCl₂, pH 6.8 adjusted with KOH) ^70,71^. Azide-labeled tubulin was prepared by reacting tubulin with N₃-PEG₄-NHS, following the standard protocol used for fluorescent dye conjugation ^49,72^. Tubulin concentration was determined from absorbance at 280 nm using a UV spectrophotometer (Nanodrop 2000c). Recombinant kinesin-1, consisting of the first 573 amino acids of human kinesin-1, was prepared as previously described ^51^.

### Complementary DNA Sequences for MT Modification

Two complementary DNA strands were used to modify MTs: 5′-TTGTTGTTGTTGTTGTTGTTGTTG-3′ (DNA1) and 5′-CAACAACAACAACAACAACAACAA-3′ (DNA2) (Table S1). 5′-dibenzo-cyclooctyne (DBCO) modified DNAs were purchased from Hokkaido System Science Co., Ltd.. DNA quality was confirmed by polyacrylamide gel electrophoresis, and sequences were verified by liquid chromatography–electrospray ionization mass spectrometry. Extinction coefficients were determined from calibration curves of absorbance versus concentration.

### Preparation of MTs

MTs were polymerized from a mixture of dye-labeled and azide-labeled tubulin (molar ratio 1:4) in polymerization buffer (80 mM PIPES, 1 mM EGTA, 1 mM MgCl₂, 1 mM GTP, pH ∼6.8), to a final tubulin concentration of 56 μM. For efficient polymerization, DMSO was added to 5% (v/v), and polymerization proceeded for 30 min at 37 °C. Immediately after, MTs were stabilized with 1 μL of 4× BRB80 and 0.5 μL of 1 mM taxol (in DMSO).

DNA modification was performed using copper-free azide–alkyne cycloaddition. Briefly, 3.5 μL of DBCO-DNA (200 μM) was added to 10 μL of azide-MTs (56 μM) and incubated for 6 h at 37 °C ^73^. DNA-modified MTs were purified by centrifugation through cushion buffer (BRB80 + 60% glycerol) at 54,000 rpm (19500 × g) for 1 h at 37 °C. Pellets were washed once with BRB80 supplemented with 1 mM taxol (BRB80P) and suspended in 15 μL BRB80P.

### Measurement of the labeling ratio of DNA to tubulin

DNA-modified MTs were depolymerized on ice overnight, yielding DNA-conjugated tubulin dimers. Absorbance spectra were measured (Nanodrop™ 2000c, Thermo Fisher Scientific Inc.) and deconvoluted by Gaussian fitting at 260 nm and 280 nm. Concentrations of DNA and tubulin were calculated using the Beer–Lambert law and their molar extinction coefficients. The DNA-to-tubulin labeling ratios were determined as 93% (DNA1) and 102% (DNA2), respectively (Table S1) ^49^.

### Preparation of flow cell and motility assays for swarm rings

Flow cells were constructed from two coverslips (MATSUNAMI, Inc.) separated by a parafilm spacer (29 × 29 mm² with a 22 × 1.5 mm² channel), producing an internal channel of ∼18 × 1.5 × 0.45 mm³.

Flow cells were first filled and incubated with 5 μL casein buffer (BRB80 supplemented with 0.5 mg/mL casein) for 3 min, followed by 5 μL kinesin solution (400 nM) for 5 min. The flow cell was washed with 10 μL wash buffer (BRB80 supplemented with 1 mM DTT,

0.5 mg/mL casein, 4.5 mg/mL glucose, 50 U/mL glucose oxidase, 50 U/mL catalase, 10 μM taxol). DNA-modified MTs were sequentially introduced: 5 μL of DNA1-modified MTs was incubated for 2 min, rinsed, then 5 μL of DNA2-modified MTs was incubated for 2 min. After washing, motility was initiated by introducing ATP buffer (wash buffer supplemented with 5 mM ATP and 0.2% methylcellulose) ^28^. The time of ATP addition was set as *t* = 0.

MT swarms spontaneously assembled into circulating rings within 30–40 min. The association ratio of red and green MTs, *A(t),* was determined by counting single MTs (not associated into swarm) at time *t, N*(*t*) relative to the initial number *N*(0) at *t* = 0., as:

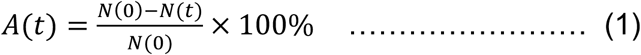

The mean association ratio was obtained from four regions of interest (2500 μm² each).

### Modification of magnetic beads with DNA

Superparamagnetic Dynabeads™ Amine (M-270) (Thermo Fisher Scientific, 2.8 μm diameter) were used for force determination experiments. These beads consist of a polystyrene matrix embedded with iron oxide nanoparticles and are coated with a hydrophilic glycidyl ether layer that seals the magnetic core. The bead surface is functionalized with primary amine groups (∼1–4 μmol/g), enabling covalent coupling to biomolecules. The beads exhibit superparamagnetic behavior, ensuring reversible magnetization without remanence, minimal aggregation in aqueous buffers, and autofluorescence under excitation.

To allow tethering to DNA-modified MTs, beads were functionalized with DNA2 (complementary to DNA1) via standard crosslinking chemistry.

### Attachment of beads to swarm rings

After swarm ring formation (∼20–30 min following ATP addition), DNA2 labeled Dynabeads (0.13 µg each, dispersed in ATP buffer) were introduced into the flow cell. Beads attached selectively to DNA1 modified MTs through complementary hybridization of DNA1 and DNA2 in a zipping geometry (Fig. S3). Once tethered, the beads were carried along the circular swarm propelled by multiple kinesin motors.

For fluorescence observation, MT swarms were assembled from DNA1 modified red MTs and DNA2 modified green MTs. After swarm formation, the individual colors could not be distinguished; spectral overlap of TRITC and GFP channels caused the swarms to appear yellow. Bead loading and transport were verified by time-lapse fluorescence imaging (Fig. S4).

Because swarms formed on both the upper and lower surfaces of the flow cell, but the beads sedimented under gravity, loading occurred predominantly at the lower surface where beads came into close contact with MT swarms.

### Custom-built electromagnetic tweezer (EMTw) system

A custom electromagnetic EMTw was designed in Autodesk Inventor, constructed accordingly, and integrated with a fluorescence microscope for simultaneous magnetic force application and imaging (Figure 2A, Fig. S6). The electromagnet consisted of a solenoid coil (6500 turns of 0.15 mm enamel-coated copper wire, resistance 240 Ω) wound around an MC Nylon® shell, enclosing a cylindrical S10C steel core with a tapered, exchangeable tip. The S10C material (0.08–0.13% carbon) provided high conductivity and low tensile strength, suitable for stable magnetic field generation.

The EMTw body was mounted on precision micro-actuators for height and lateral positioning, and a micro-rotary actuator enabled angular adjustment of the tapered tip for directional force application. The system was powered by a regulated DC supply (KIKUSUI PMX70-1A), capable of delivering 0.001–0.150 A at up to 36 V where the coil resistance is 240 Ω. Force magnitude scaled with applied current until saturation of the core material. At higher currents, resistive heating reduced magnetic efficiency; therefore, voltage and current settings were optimized to minimize thermal effects.

### Force calibration of Dynabeads using EMTw

Prior to swarm experiments, force calibration of Dynabeads was performed in 80% (v/v) glycerol (viscosity η = 0.0737 Pa·s at 25 °C) to reduce bead motion and counteract sedimentation. Beads were introduced into a flow cell, and the EMTw tip was positioned ∼200 µm from the observation region to apply lateral magnetic forces. Under applied EMF, bead trajectories were recorded using time-lapse fluorescence microscopy.

Displacement was obtained from trajectory analysis, and velocities were used to calculate the applied force (F) via Stokes’ law (Eq. i, main text), as shown in Figure 2B ^53^. At fixed distances, force scaled linearly with applied voltage (*F* ∝ 𝑣), providing straightforward calibration of EMTw force output.

### Application of EMF to the bead-attached swarm by EMTw

During EMF applcation, bead-loaded swarms were positioned within the microscope field of view at distances of 200–300 μm from the EMTw tip. At this range, applied EMF remained stable against small (̴ 5 um) distance fluctuations. Applied EMF was increased in stepwise fashion, while swarm dynamics were monitored by fluorescence microscopy. The alternating application of EMF and rest intervals (no EMF) allowed assessment of swarm velocity modulation in response to external force.

### Landing rate experiment for kinesin surface density

The density of kinesins on the glass surface was estimated by a landing rate assay ^56,57^. ATTO565-labeled GTP-MTs (average length 8.98 ± 5.16 μm (SD)) were introduced into flow cells coated with kinesin at 400 nM, matching the concentration used in swarm experiments. Kinesin stock solution was serially diluted (dilution factors *ξ*, Table S2), and MT landings on the surface were recorded by time-lapse fluorescence imaging for 10 min. The cumulative number of landed MTs, *N*(*t*), was plotted in Fig. S12 and fitted with the following equation:

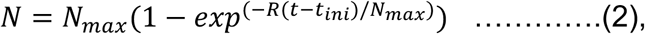

where *N* is the number of MTs, *N*_max_ is the maximum number of landed MTs for each dilution, *R* is the landing rate, *t* is time, and *t*_ini_ = 120 s accounts for the lag between MT addition and image acquisition. *N*_max_ and *R* are the fitting parameters.

The landing rate *R* for each kinesin concentration was obtained from the fitted curves in Fig. S12 (Eq. 2). The resulting *R* values were then plotted against the dilution factor *ξ* (Fig. S13) and fitted using the following equation to estimate the kinesin surface density:

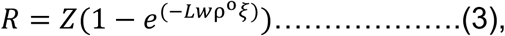

where *Z* is a constant, *ξ* is the dilution factor, *L* and *w* denote the average MT length and width (20 nm), and *ρ*° represents the kinesin surface density corresponding to the 400 nM stock solution. From multiple trials (*n* = 8), the average kinesin density, *ρ*° was determined to be ∼ 61.05 ± 22.04 µm^-2^.

### Analysis of trajectories, velocity, and force evaluation

The trajectories of bead-attached MT swarm rings, with and without applied EMF, were extracted from fluorescence image sequences using ImageJ (NIH). Instantaneous velocities were calculated from frame-to-frame displacements, and mean velocities were obtained over defined time intervals. The influence of EMF on swarm dynamics was assessed by comparing trajectory maps and velocity profiles across alternating force application cycles. A total of 26 complete events were obtained for quantitative force analysis.

### Estimation of parameters and statistical analysis

Fluorescence images were analyzed using NIS-Elements AR (version 5.30.02, Nikon). Quantitative analysis of swarm dimensions (diameter, width, annulus area) and fluorescence intensity profiles was performed using this software. Statistical analysis and plotting were carried out with OriginPro 2019 (OriginLab, USA). Reported values represent mean ± standard deviation (SD) unless otherwise stated. Sample sizes are provided in figure captions.

## Acknowledgments

We acknowledge A. Konagaya, Professor, Tokyo Institute of Technology; Kazuhiro Oiwa, Professor, National Institute of Information and Communications Technology, Japan; Stefan Diez, professor, Max Plank Institute of Molecular Cell Biology and Genetics, Germany; Jakia Jannat Keya, JSPS Postdoctoral researcher, Hokkaido University; and S.R. Nasrin, JSPS Postdoctoral researcher, Kyoto University for their valuable time and suggestions to improve our work.

## Author contributions

Conceptualization: A.K.

Methodology: M.R.R., M.A.

Investigation: M.R.R.

Visualization: M.R.R.

Funding acquisition: A.K., A.M.R.K., and T.H.

Project administration: M.R.R, M.A., A.M.R.K., K.S., and A.K.

Supervision: A.K.

Writing: M.R.R

Writing—review and editing: M.R.R., M.A., A.M.R.K., A.

Kuzuya, T.H., K.S., I.K., M.T., and A.K.

## Supporting Information

**Figure S1.**
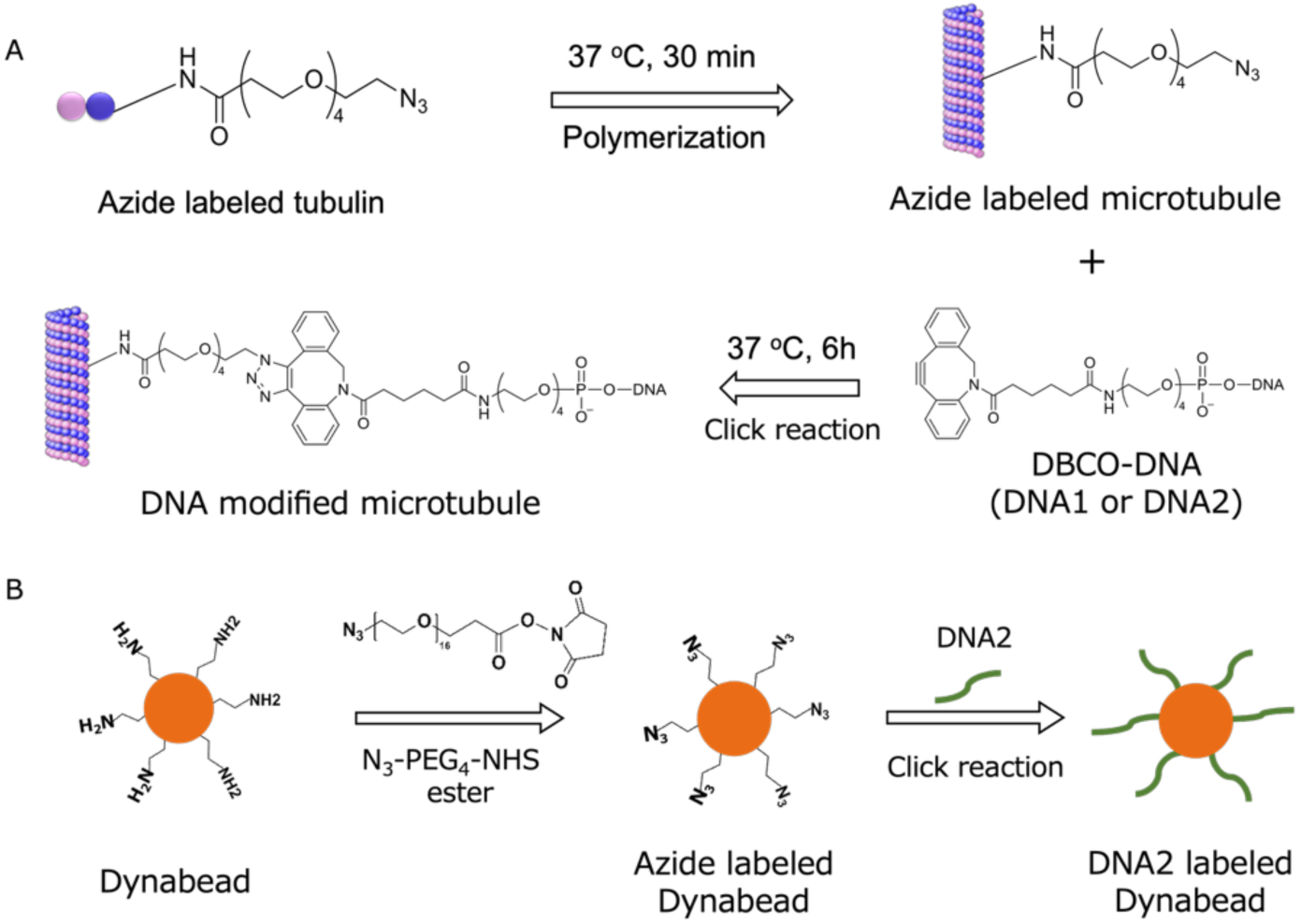
Preparation of DNA-modified microtubules (MTs/swarm units) and DNA-modified beads. (A) Schematic representation of the preparation of MT swarm units using complementary DNA sequences. Microtubules were polymerized from azide-functionalized tubulin mixed with ATTO550- or ATTO488-labeled tubulin (red and green MTs, respectively) at a 1:4 ratio in the presence of GTP at 37 °C. Each set of MTs was then conjugated with complementary DNAs (DNA1 and DNA2; Fig. 1C.i) via copper-free click chemistry, in which the DBCO group at the 5′ end of the DNA reacts with the azide groups on the MT surface. (B) Schematic representation of the DNA modification of Dynabead.

**Figure S2.**
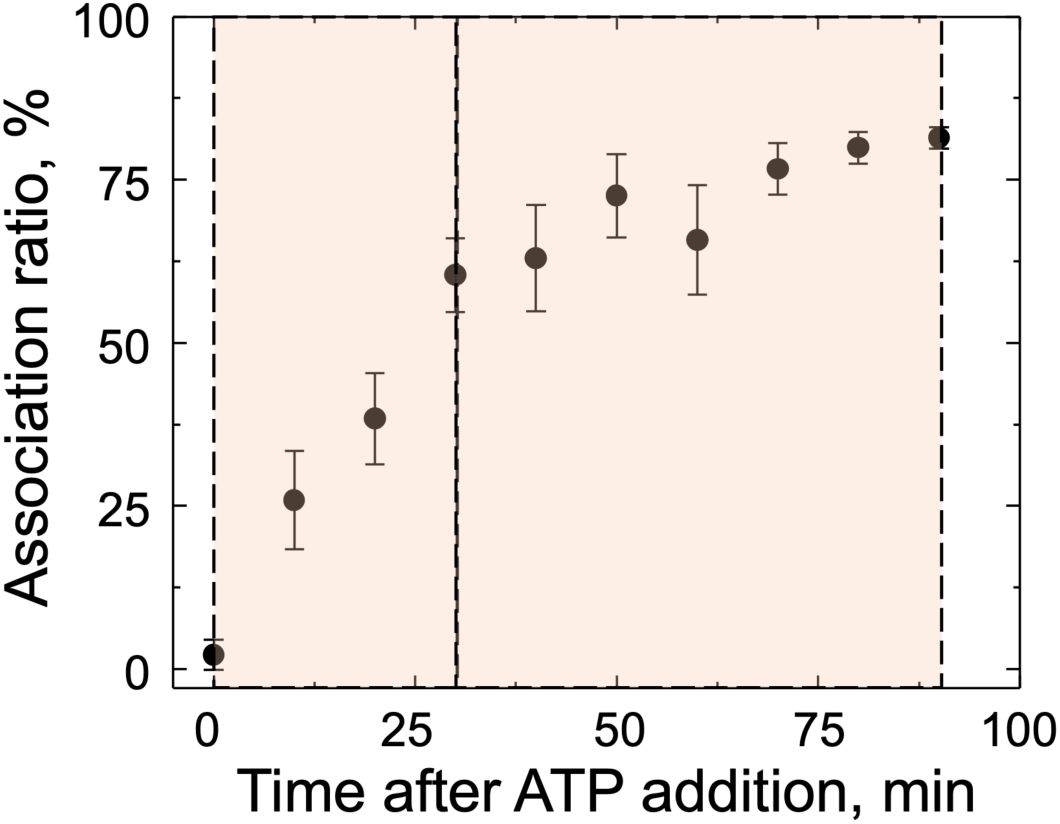
Time-dependent association; The swarm rings of MTs grown rapidly up to 30 min and then gradually reached a plateau within 90 min. The association ratio at time t was calculated as, 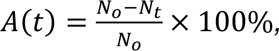 where *N*_*o*_ and *N*_*t*_ represent the numbers of single MTs at the initial time t_o_, and at time t, respectively.

**Figure S3.**
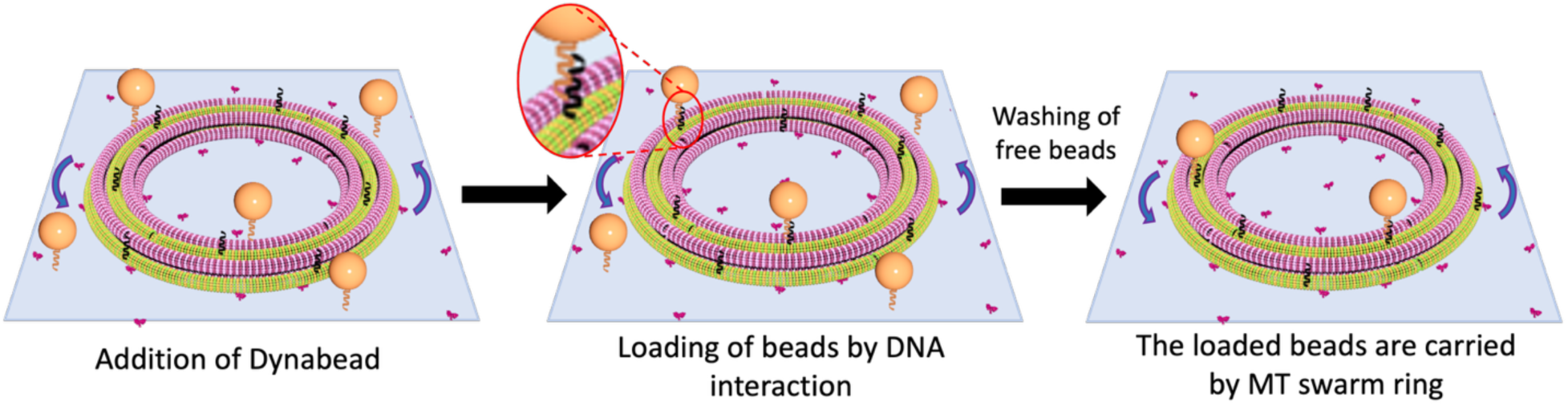
Schematic representation of bead attachment to a MT swarm via DNA zipping. The DNA strands on the bead and MT swarm hybridize through complementary base pairing (DNA zipping), enabling the bead to attach to the swarm ring. Once attached, the bead is carried by the kinesin propelled swarm along a circular trajectory. The schematic is not drawn to scale.

**Figure S4.**
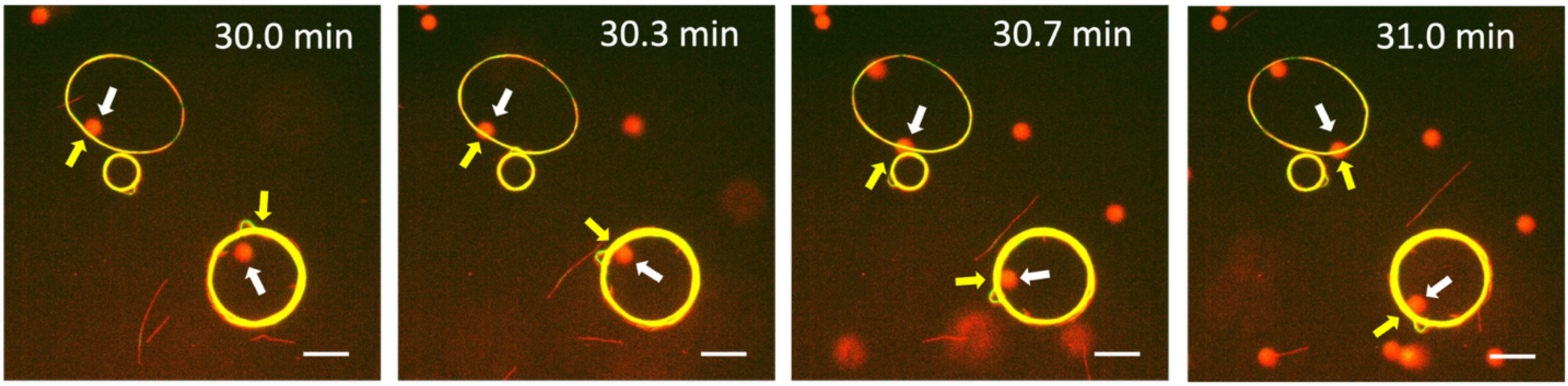
Time-lapse fluorescence microscopy images of DNA-assisted bead loading by a MT swarm ring. Representative fluorescence images show beads (DNA2-modified) attaching to MTs (DNA1-modified red MTs) and subsequently transported along the circular path of the swarm. Red and green MTs were visualized under TRITC and GFP filters, respectively, appearing together as yellow due to overlap. Scale bar = 10 μm.

**Figure S5.**
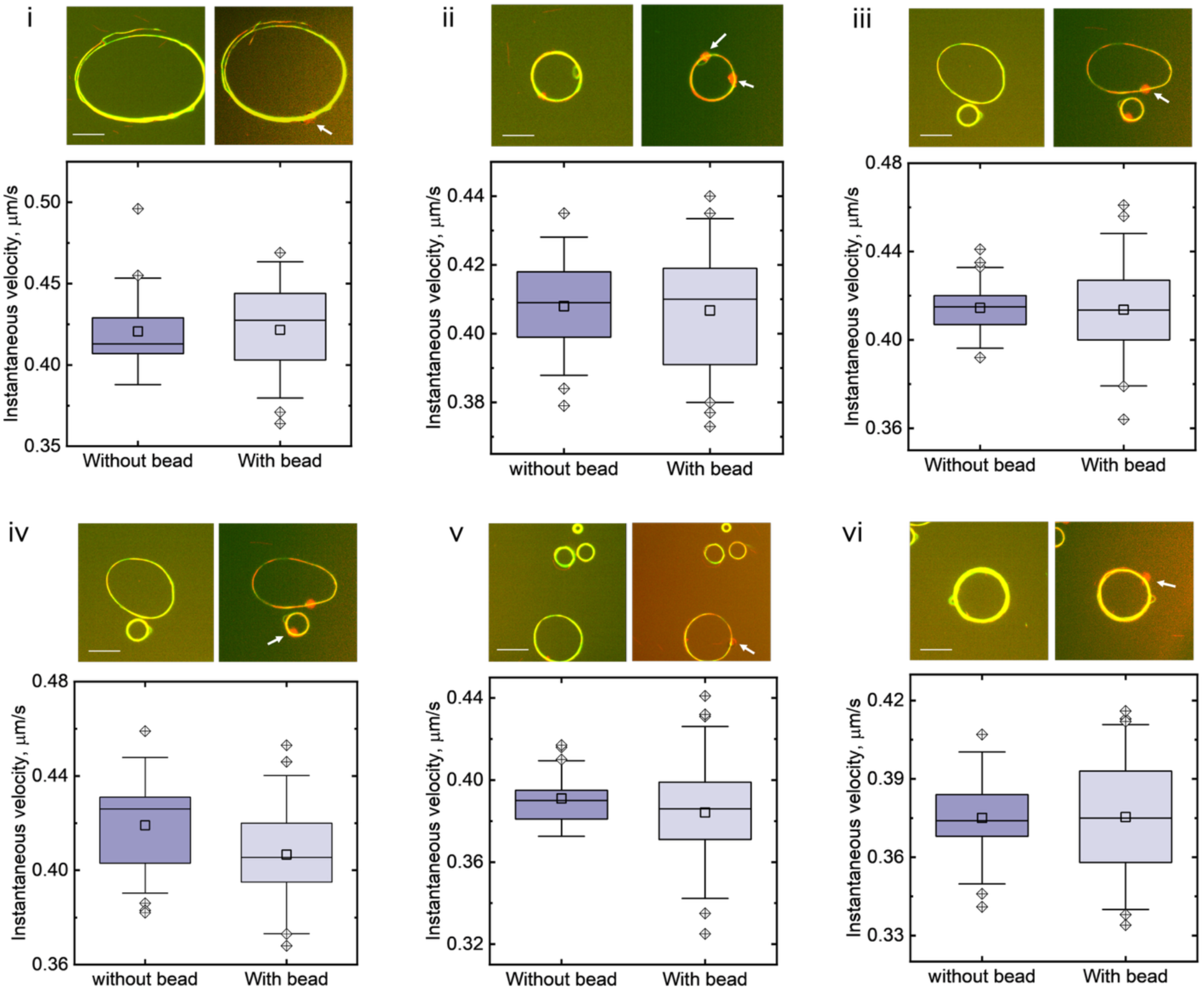
Fluorescence microscopy images and velocity analysis of kinesin-propelled MT swarms with and without bead attachment. (i–vi) Representative fluorescence images show MT swarms without beads (left) and with bead attachment (right). The accompanying box plot compares the instantaneous velocities of MT swarms with and without attached beads. Outliers and mean ± 1.5 SD are indicated. Statistical analysis using one-way ANOVA showed no significant difference between the two datasets (p > 0.01).

**Figure S6.**
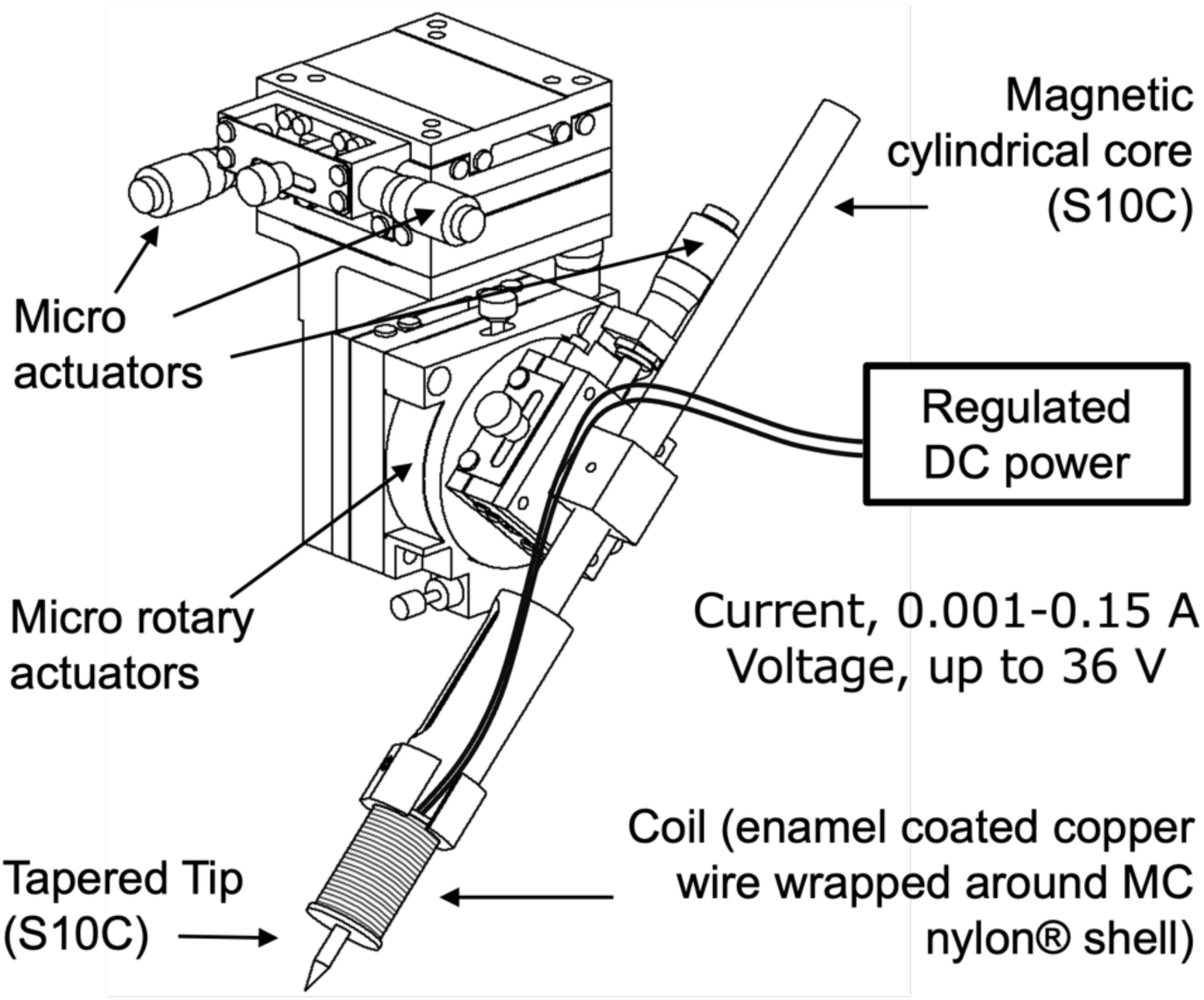
Custom-built electromagnetic tweezer (EMTw) setup. Schematic of the EMTw, modeled and assembled using Autodesk Inventor (USI), showing all components configured for laboratory operation.

### Force calibration of Dynabeads

Force–distance relationships were obtained for multiple beads under varying distances and currents and fitted with a simple power-law expression ^1^:

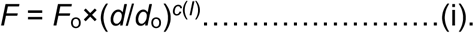

Equation (i) describes the dependence of force (*F*) on current (*I*) and distance (*d*), where *F*₀ and *d*₀ are the reference force and distance, respectively. The exponent *c(I)* represents the current-dependent distance exponent (typically *c* = –1 or –2..), characterizing the slope of the force–distance (*F–d*) curve in log–log scale.

Because current and voltage are proportional (*V* = *I*×*R*), in our setup the exponent was expressed as a voltage-dependent term, *c(V)*, allowing simplification of Equation (i) as follows:

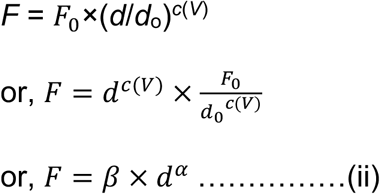

where α = *c*(*V*) and 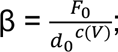 α and β are the fitting parameters, where *β* is negative (*β* < 0).

**Figure S7.**
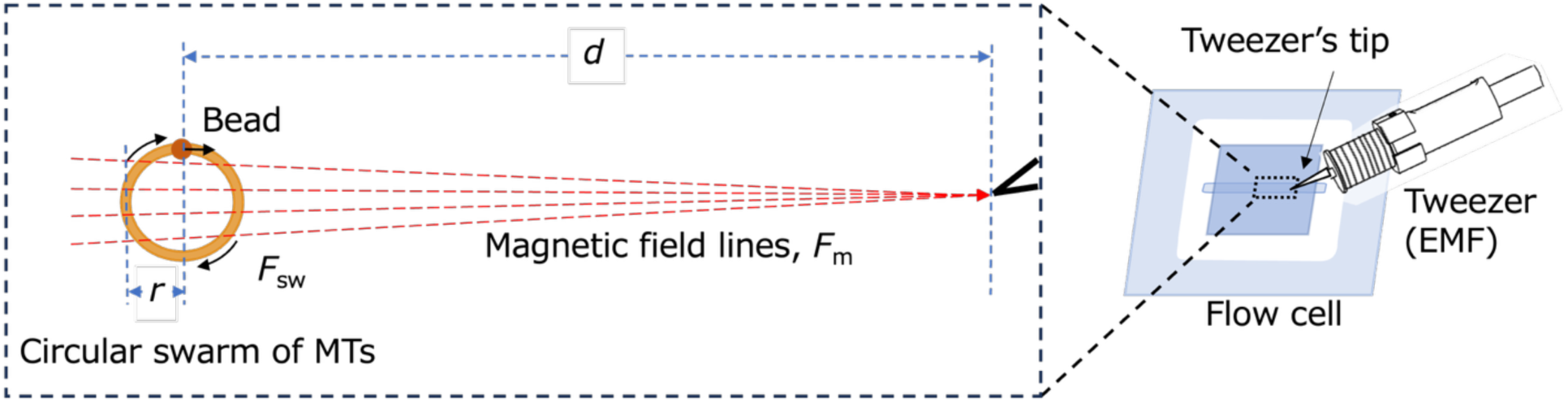
Schematic of the electromagnetic tweezer (EMTw) and bead-loaded MT swarm within the flow cell; The distance (d) between the EMTw tip and the center of the swarm is considerably larger than the swarm radius (r). The magnetic field lines are assumed to be parallel across the swarm region, with an angular deviation of approximately ±1° within the swarm area. The schematic is not drawn to scale.

**Figure S8.**
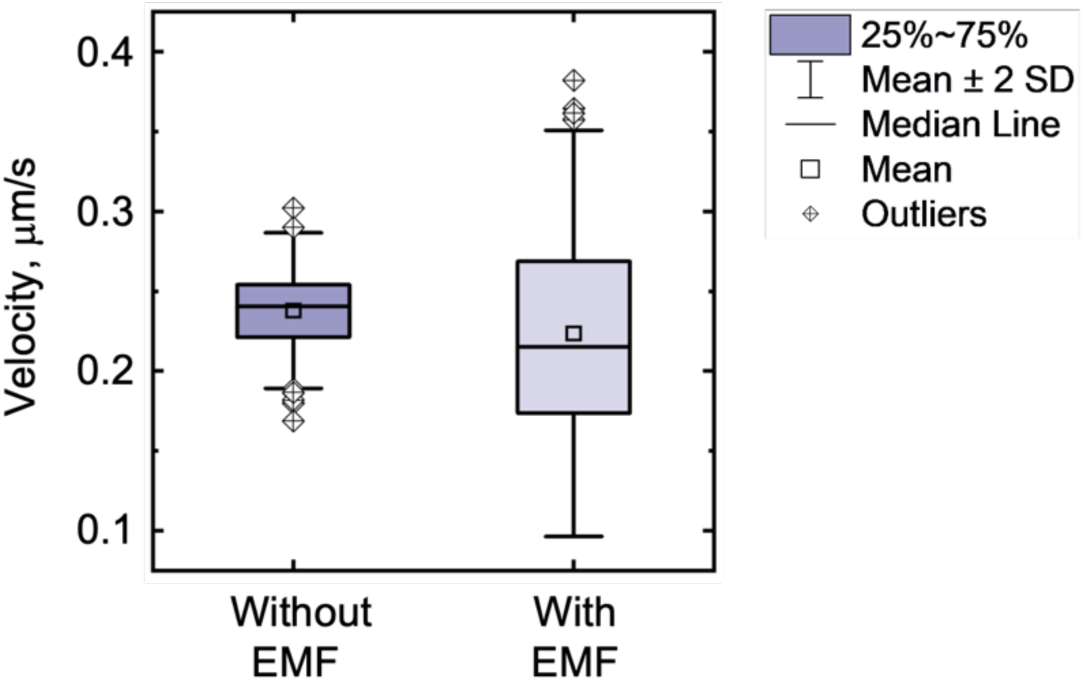
Instantaneous velocity of the MT swarm with and without applied EMF; Velocity traces were analyzed during intervals without EMF (0–300 s and 600–900 s) and with EMF applied (300–600 s and 900–1200 s). A total of n = 120 instantaneous velocity measurements were used to compute the mean values for each condition. One-way ANOVA indicated no significant difference between the mean velocities of the two conditions (p < 0.01). However, Levene’s test for equality of variances, based on squared deviations, indicated that the population variances differed significantly at the 0.01 level.

**Figure S9.**
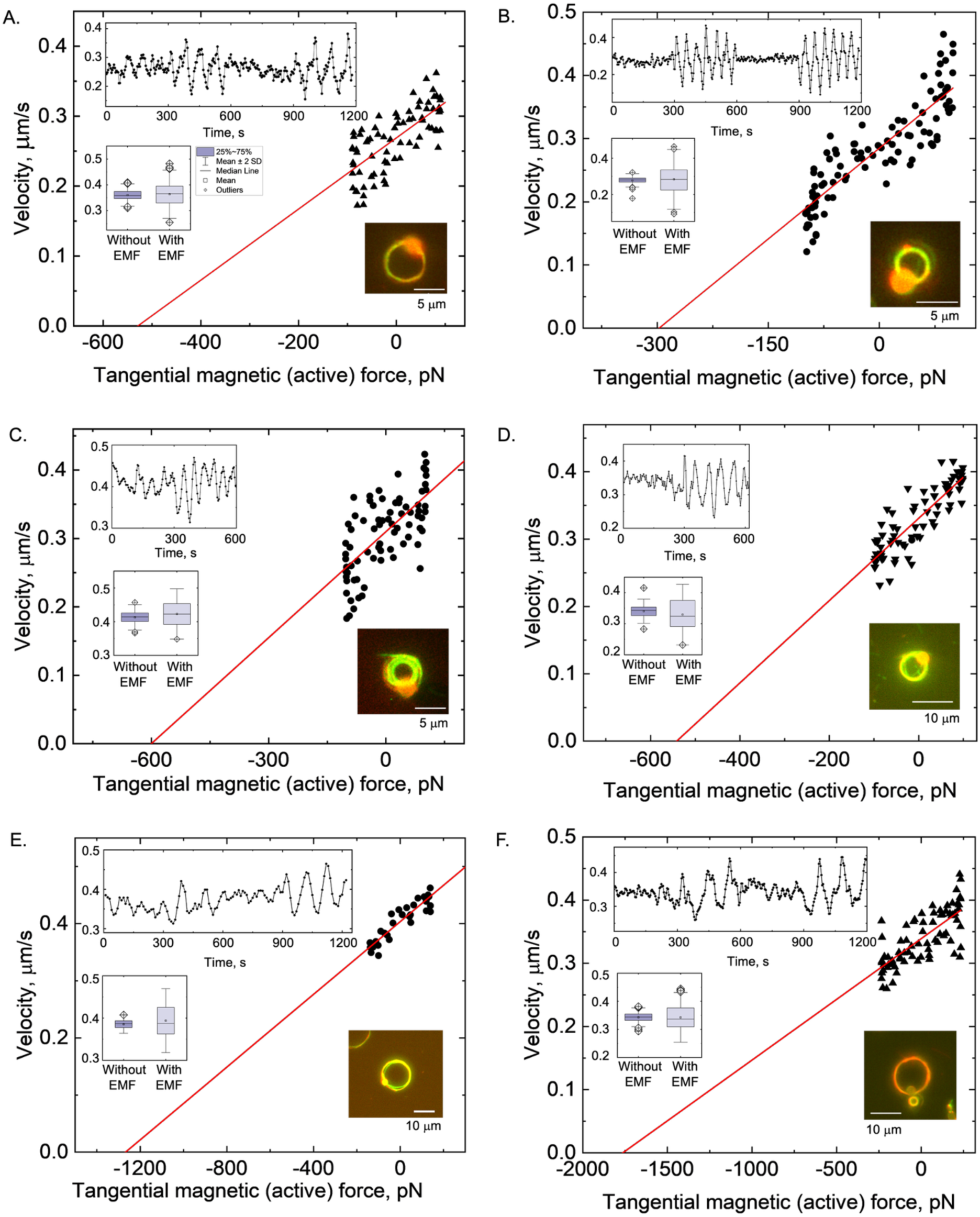
Forces generated by MT swarm rings of varying sizes. (A–F) Force measurements were obtained from multiple swarm events with different ring diameters and widths. The generated forces ranged from a few hundred to several thousand piconewtons (pN), depending on swarm size and geometry.

**Figure S10.**
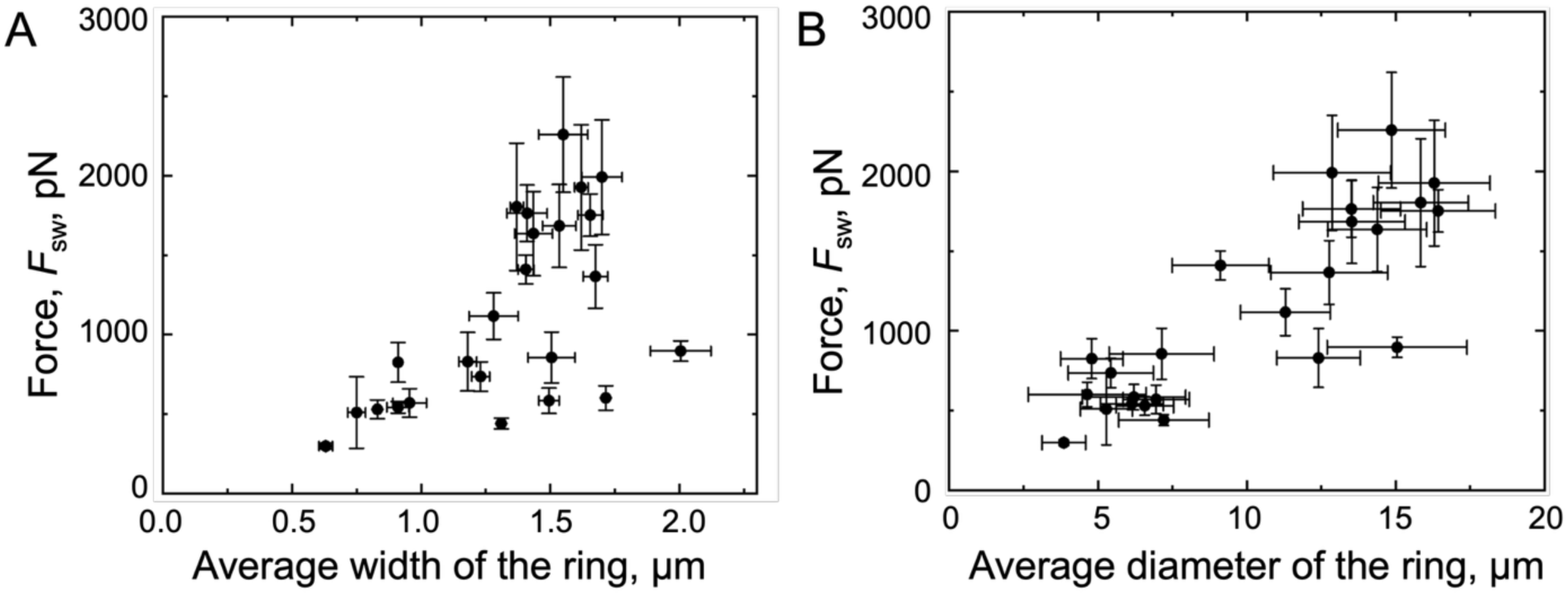
Relationship between swarm force and geometric parameters of MT rings; The force generated by the swarm F_sw_, was analyzed as a function of average ring width and diameter. Ring widths were determined from fluorescence images converted to threshold binary images. F_sw_ increased with both (A) swarm ring width, (B) average ring diameter, showing greater variability at larger widths and diameters.

**Figure S11.**
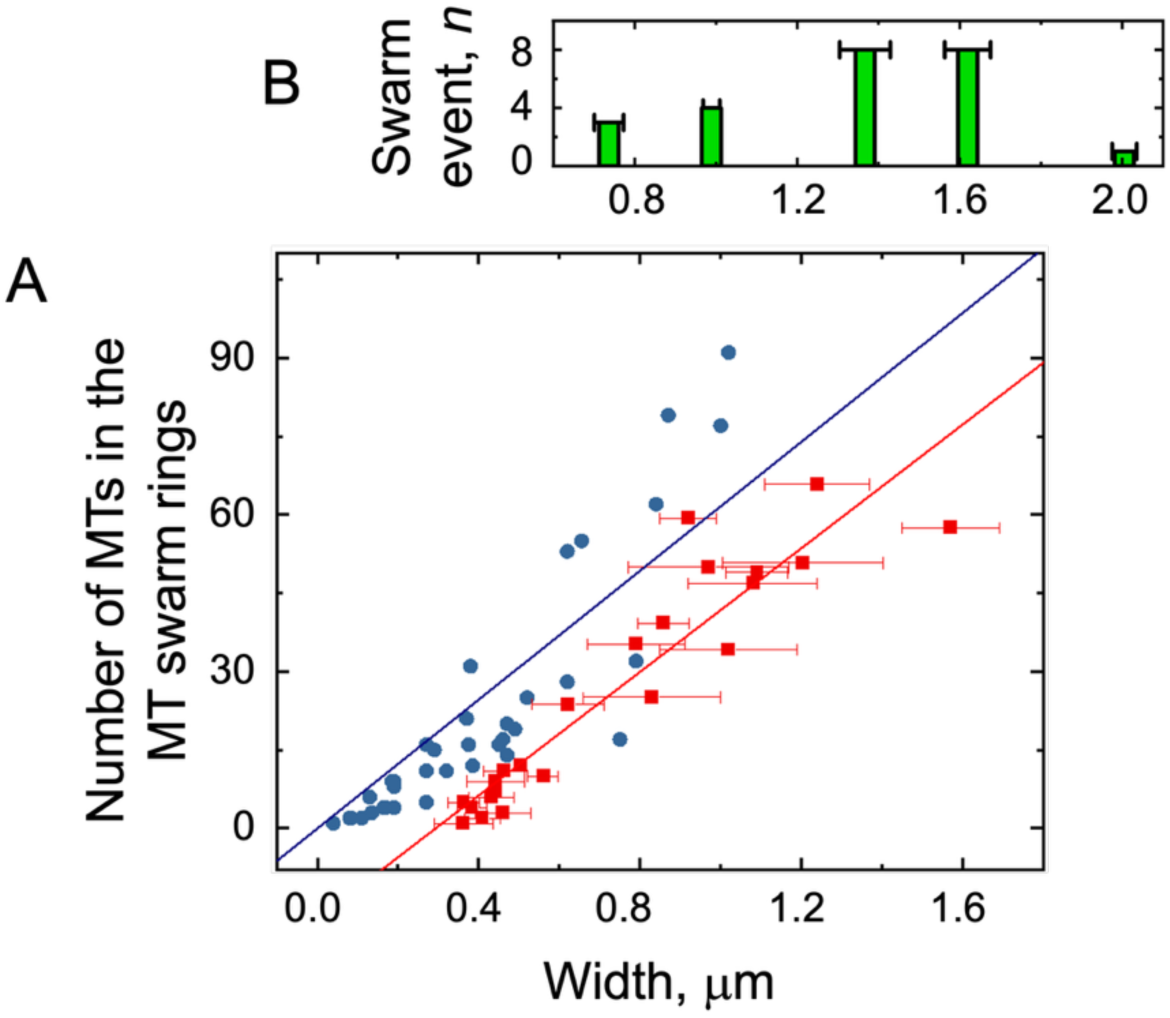
Estimation of MT numbers in swarm rings; (A) Comparison of MT counts in swarm rings obtained from photo-dissociation experiments using UV light and photo-responsive DNA, observed by fluorescence microscopy, with MT numbers estimated from height measurements using high-speed atomic force microscopy (HS-AFM) in a previous study ^2^. (B) Comparison of fluorescence widths of swarm rings used in the present force determination experiments with those reported previously. These prior studies showed that the number of MT filaments increases with ring width. In this work, swarm rings with widths greater than 1 μm were selected, consistent with a higher number of MT filaments at larger ring widths.

**Figure S12.**
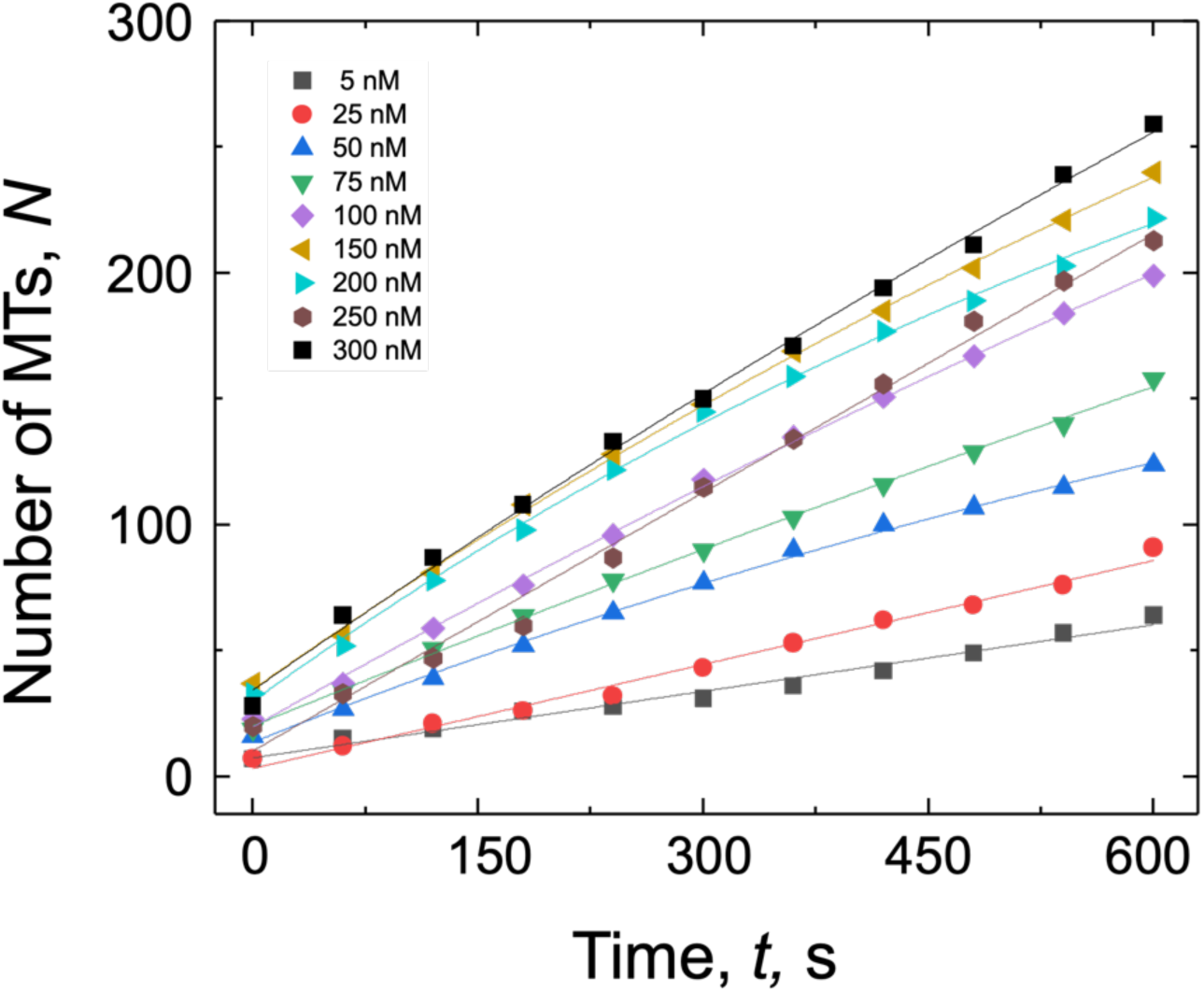
Time-dependent MT landing events at different kinesin dilutions; The number of landed MTs within the field of view was analyzed for varying dilution factors of kinesin. Data were fitted using the landing-rate equation, *N* = *N*_*max*_(1 − *exp*^(−*R*(*t*−*tini*)/*Nmax*)^) ^3^, where N is the number of landed MTs, N_max_ is the maximum number of MTs landed for each dilution, R is the landing rate, t is time, and t_ini_ = 120 s represents the lag between MT addition and image acquisition. N_max_ and R, are fitting parameters. All fits yielded regression values (R²) greater than 0.99.

**Figure S13.**
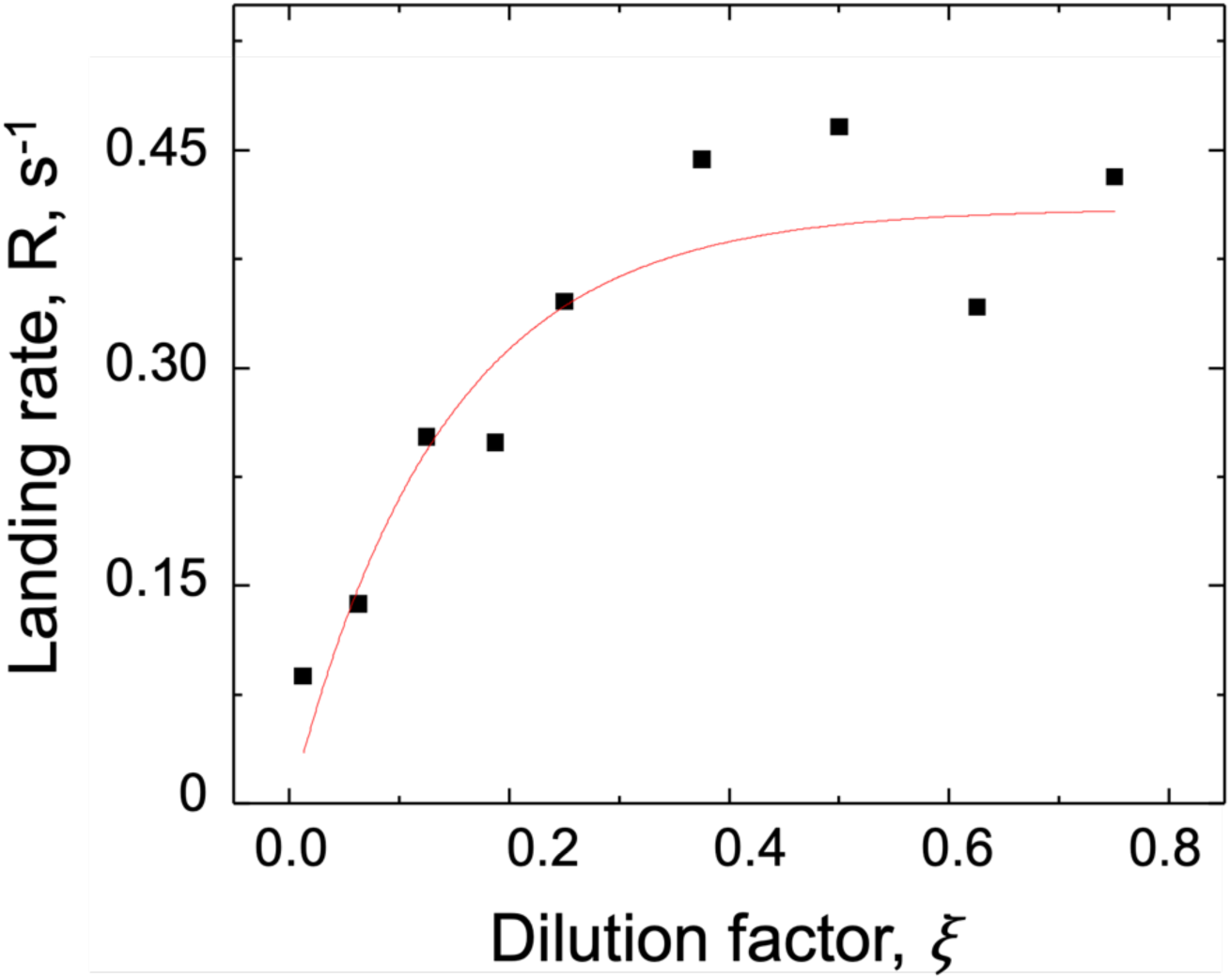
Landing rate of MTs as a function of kinesin dilution; The landing rate (R) of MTs was plotted against the dilution factor (ξ) of the 400 nM kinesin stock solution. Data were fitted using the landing-rate model: *R* = *Z*(1 − *e*^(−*Lwρoξ*)^) ^3^; where *Z* is a constant, *ξ* is the dilution factor, *ρ*^*o*^ is the kinesin surface density of the 400 nM stock, and *Lw* represents the effective MT area (*L* = average MT length; *w* = 20 nm). The fit yielded a regression coefficient (R²) of 0.85.

**Figure S14.**
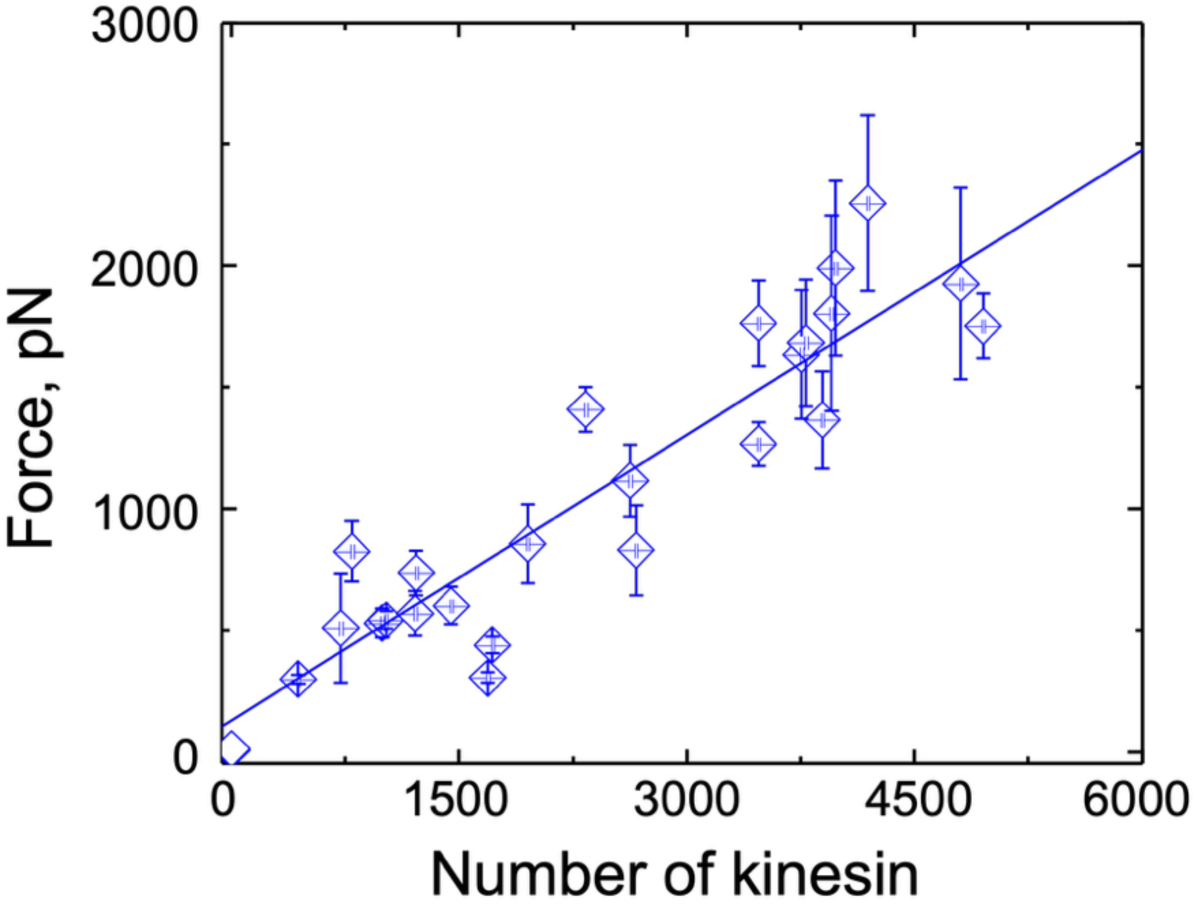
Relationship between swarm force and the number of kinesins; The force generated by each MT swarm was plotted as a function of the number of kinesins participating in the swarm, as determined from the landing-rate experiment. The swarm force increased with the number of engaged kinesins, indicating cooperative force generation.

**Figure S15.**
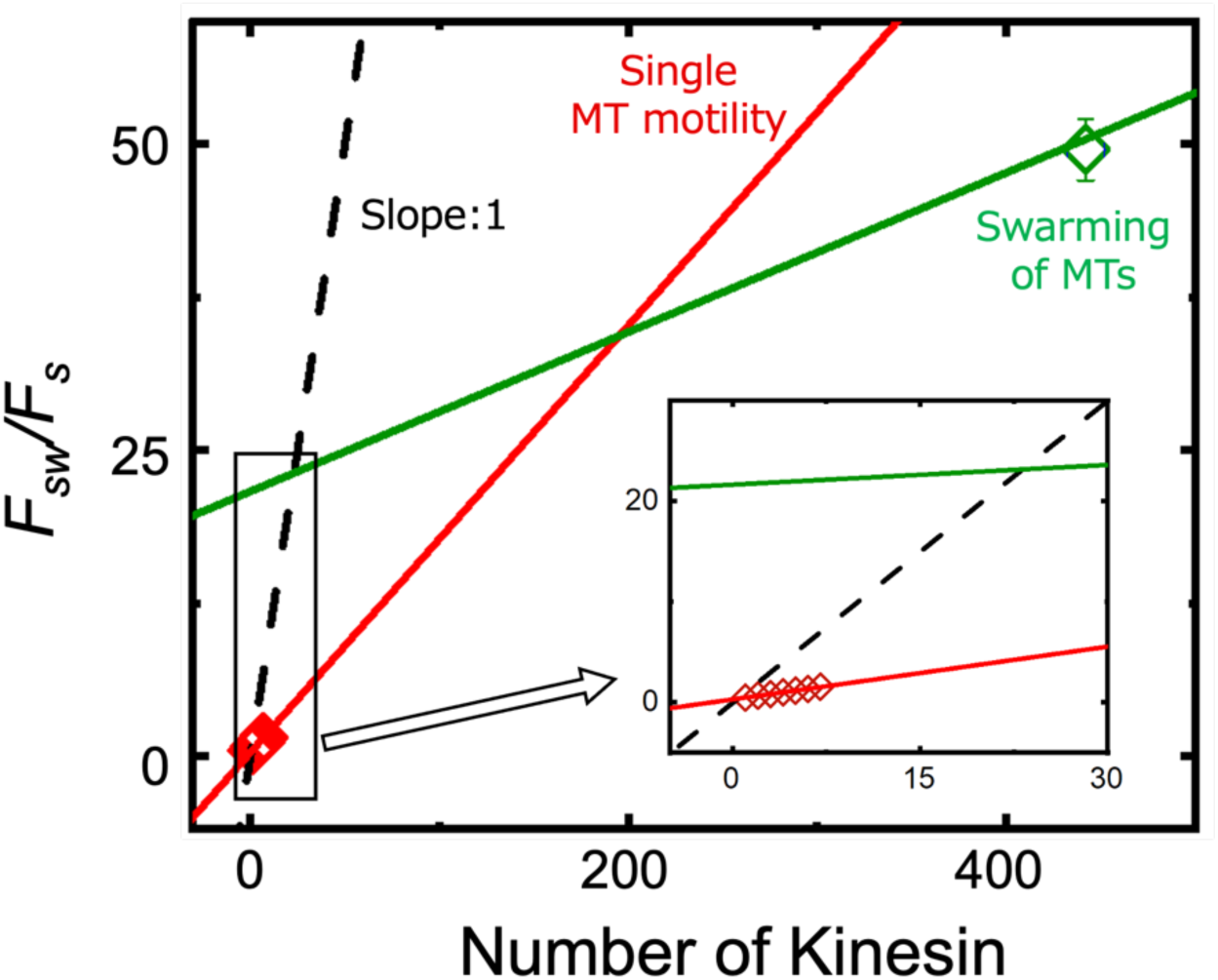
Magnified view of the lower-scale region from Figure 4B inset; The plot highlights the force generated by a few kinesins (red line), which lies in the lower range compared with the higher forces produced by the microtubule (MT) swarm (blue line). The dashed line represents a unit slope (slope = 1), corresponding to the expected additive force from individual kinesins.

**Table S1.** Labeling ratios of DNA1 and DNA2 modified MTs:

| DNA | Infeed concentration ($\mu\text{M}$) | Molar extinction coefficient ($\text{Lmol}^{-1}\text{cm}^{-1}$) | Conc. of DNA ($\mu\text{M}$) | Conc. of tubulin ($\mu\text{M}$) | DNA to Tubulin L. R. (%) |
| --- | --- | --- | --- | --- | --- |
| DNA1<br>DBCO(CAA)<sub>8</sub> | 200 | 210200 | 7.5 | 8.1 | 93.4 % |
| DNA1<br>DBCO(TTG)<sub>8</sub> | 200 | 147400 | 12.5 | 12.2 | 102.2 % |

**Table S2.** Kinesin concentrations corresponding to different dilution factors of the 400 nM stock solution.

| Kinesin concentration (nM) | Dilution factor, $\xi$ |
| --- | --- |
| 1 | 0.0025 |
| 5 | 0.0125 |
| 10 | 0.025 |
| 25 | 0.0625 |
| 50 | 0.125 |
| 75 | 0.1875 |
| 100 | 0.25 |
| 150 | 0.375 |
| 200 | 0.5 |

### Movie S1, separate file corresponds to Figure 3

Time-lapse fluorescence movie of the bead attached-swarm ring without applying EMF (0 V, from 0 to 300 s and 600 to 900 s) and with applying EMF (10 V, from 300 to 600 s and 900 to 1200 s). The event corresponding to the movie is presented in Figure 3 in the main manuscript. The velocity was controlled by applying EMF using EMTw, to the single bead-attached circular swarm to determine the swarm force. Scale bar 5 µm. The movie is 100 times faster than the original speed.

### Movie S2, separate file corresponds to Fig. S9(A-F)

Time-lapse fluorescence movie of the bead-attached swarm rings alternating conditions of (a) no electric field (0 V; 0-300 s and 600-900 s) and applied EMF (10 V; 300- 600 s and 900-1200 s), (b) no electric field applied during 0- 300 s and 700- 900 s) and with applying EMF of 10 V during 300-800 s and 900-1200 s, (c) no EMF (0 V; from 0- 300) and with applying EMF (18 V; from 300-600 s), (d) no EMF (0 V, from 0-300 s) and with applying EMF (10 V; from 300-600 s), (e) without applying EMF (0 V; from 0-300 s and 600-900 s) and with applying EMF (15 V, from 300-600 s and 900-1200 s), (f) without applying EMF (0 V, from 0-300 s and 600-900 s) and with applying EMF (18 V; from 300- 600 s and 900-1200 s). The events corresponding to the movies are presented in Fig. S9 in the supporting information. The velocity was controlled by applying EMF using EMTw, to the single bead-attached circular swarm to determine the swarm force. Scale bar 5 µm.

## Reference

(1) Whitesides, G. M.; Grzybowski, B. Self-Assembly at All Scales. Science 2002, 295 (5564), 2418–2421.

(2) Beshers, S. N.; Fewell, J. H. Models of Division of Labor in Social Insects. Annu. Rev. Entomol. 2001, 46 (1), 413–440.

(3) Niven, J. E. How Honeybees Break a Decision-Making Deadlock. Science 2012, 335 (6064), 43–44.

(4) Feinerman, O.; Pinkoviezky, I.; Gelblum, A.; Fonio, E.; Gov, N. S. The Physics of Cooperative Transport in Groups of Ants. Nat. Phys. 2018, 14 (7), 683–693. 10.1038/s41567-018-0107-y.

(5) Mlot, N. J.; Tovey, C. A.; Hu, D. L. Fire Ants Self-Assemble into Waterproof Rafts to Survive Floods. Proc. Natl. Acad. Sci. U. S. A. 2011, 108 (19), 7669–7673. 10.1073/pnas.1016658108.

(6) Shklarsh, A.; Finkelshtein, A.; Ariel, G.; Kalisman, O.; Ingham, C.; Ben-Jacob, E. Collective Navigation of Cargo-Carrying Swarms. Interface Focus 2012, 2 (6), 786–798. 10.1098/rsfs.2012.0029.

(7) Gelblum, A.; Pinkoviezky, I.; Fonio, E.; Ghosh, A.; Gov, N.; Feinerman, O. Ant Groups Optimally Amplify the Effect of Transiently Informed Individuals. Nat. Commun. 2015, 6. 10.1038/ncomms8729.

(8) Turgut, A. E.; Çelikkanat, H.; Gökçe, F.; Şahin, E. Self-Organized Flocking in Mobile Robot Swarms. Swarm Intell. 2008, 2 (2), 97–120. 10.1007/s11721-008-0016-2.

(9) Wei, H.; Chen, Y.; Tan, J.; Wang, T. Sambot: A Self-Assembly Modular Robot System. IEEEASME Trans. Mechatron. 2011, 16 (4), 745–757. 10.1109/TMECH.2010.2085009.

(10) Wang, W.; Duan, W.; Ahmed, S.; Sen, A.; Mallouk, T. E. From One to Many: Dynamic Assembly and Collective Behavior of Self-Propelled Colloidal Motors. Acc. Chem. Res. 2015, 48 (7), 1938–1946.

(11) Rubenstein, M.; Cornejo, A.; Nagpal, R. Programmable Self-Assembly in a Thousand-Robot Swarm. Science 2014, 345 (6198), 795–799. 10.1126/science.1254295.

(12) Palagi, S.; Fischer, P. Bioinspired Microrobots. Nat. Rev. Mater. 2018, 3 (6), 113–124. 10.1038/s41578-018-0016-9.

(13) Dorigo, M.; Théraulaz, G.; Trianni, V.; Dorigo, M.; Théraulaz, G.; Trianni, V. Reflections on the Future of Swarm Robotics. *Sci*. Robot. 2021, 5 (49).

(14) Vale, R. D. The Molecular Motor Toolbox for Intracellular Transport. Cell 2003, 112 (4), 467–480. 10.1016/s0092-8674(03)00111-9.

(15) Sato, Y.; Hiratsuka, Y.; Kawamata, I.; Murata, S.; Nomura, S. I. M. Micrometer-Sized Molecular Robot Changes Its Shape in Response to Signal Molecules. Sci. Robot. 2017, 2 (4).

(16) Keya, J. J.; Kabir, A. M. R.; Inoue, D.; Sada, K.; Hess, H.; Kuzuya, A.; Kakugo, A. Control of Swarming of Molecular Robots. Sci. Rep. 2018, 8 (1), 1–10.

(17) Dumont, E. L. P.; Do, C.; Hess, H. Molecular Wear of Microtubules Propelled by Surface-Adhered Kinesins. Nat. Nanotechnol. 2015, 10 (2), 166–169.

(18) Clemmens, J.; Hess, H.; Doot, R.; Matzke, C. M.; Bachand, G. D.; Vogel, V. Motor-Protein “Roundabouts”: Microtubules Moving on Kinesin-Coated Tracks through Engineered Networks. Lab. Chip 2004, 4 (2), 83–86.

(19) Bachand, G. D.; Rivera, S. B.; Boal, A. K.; Gaudioso, J.; Liu, J.; Bunker, B. C. Assembly and Transport of Nanocrystal CdSe Quantum Dot Nanocomposites Using Microtubules and Kinesin Motor Proteins. Nano Lett. 2004, 4 (5), 817–821.

(20) Roberts, A. J.; Kon, T.; Knight, P. J.; Sutoh, K.; Burgess, S. A. Functions and Mechanics of Dynein Motor Proteins. Nat. Rev. Mol. Cell Biol. 2013, 14 (11), 713–726. 10.1038/nrm3667.

(21) Huber, L.; Suzuki, R.; Krüger, T.; Frey, E.; Bausch, A. R. Emergence of Coexisting Ordered States in Active Matter Systems. Science 2018, 361 (6399), 255–258. 10.1126/science.aao5434.

(22) Keya, J. J.; Suzuki, R.; Kabir, A. M. R.; Inoue, D.; Asanuma, H.; Sada, K.; Hess, H.; Kuzuya, A.; Kakugo, A. DNA-Assisted Swarm Control in a Biomolecular Motor System. Nat. Commun. 2018, 9 (1), 4–11.

(23) Hess, H.; Clemmens, J.; Brunner, C.; Doot, R.; Luna, S.; Ernst, K. H.; Vogel, V. Molecular Self-Assembly of “Nanowires” and “Nanospools” Using Active Transport. Nano Lett. 2005, 5 (4), 629–633.

(24) Hess, H.; Ross, J. L. Non-Equilibrium Assembly of Microtubules: From Molecules to Autonomous Chemical Robots. Chem. Soc. Rev. 2017, 46 (18), 5570–5587.

(25) Gardner, M. K.; Charlebois, B. D.; Jánosi, I. M.; Howard, J.; Hunt, A. J.; Odde, D. J. Rapid Microtubule Self-Assembly Kinetics. Cell 2011, 146 (4), 582–592.

(26) Hess, H. Self-Assembly Driven by Molecular Motors. Soft Matter 2006, 2 (8), 669–677.

(27) Ishii, S.; Akter, M.; Murayama, K.; Kabir, A. M. R.; Asanuma, H.; Sada, K.; Kakugo, A. Kinesin Motors Driven Microtubule Swarming Triggered by UV Light. Polym. J. 2022, 54 (12), 1501–1507.

(28) Akter, M.; Keya, J. J.; Kayano, K.; Kabir, A. M. R.; Inoue, D.; Hess, H.; Sada, K.; Kuzuya, A.; Hiroyuki, A.; Kakugo, A. Cooperative Cargo Transportation by a Swarm of Molecular Machines. *Sci*. Robot. 2022, 7 (65), eabm0677.

(29) Kawamata, I.; Nishiyama, K.; Matsumoto, D.; Ichiseki, S.; Keya, J. J.; Okuyama, K.; Ichikawa, M.; Kabir, A. R.; Sato, Y.; Inoue, D.; Nomura, M. Autonomous Assembly and Disassembly of Gliding Molecular Robots Regulated by a DNA- - Based Molecular Controller. Sci. Adv. 2024, 10 (eadn4490), 1–9.

(30) Ariga, T.; Tomishige, M.; Mizuno, D. Nonequilibrium Energetics of Molecular Motor Kinesin. Phys. Rev. Lett. 2018, 121 (21), 218101.

(31) Bustamante, C. J.; Chemla, Y. R.; Liu, S.; Wang, M. D. Optical Tweezers in Single-Molecule Biophysics. Nat. Rev. Methods Primer 2021, 1 (1).

(32) Svoboda, K.; Block, S. M. Force and Velocity Measured for Single Kinesin Molecules. Cell 1994, 77 (5), 773–784.

(33) Block, S. M.; Asbury, C. L.; Shaevitz, J. W.; Lang, M. J. Probing the Kinesin Reaction Cycle with a 2D Optical Force Clamp. Proc. Natl. Acad. Sci. 2003, 100 (5), 2351–2356. 10.1073/pnas.0436709100.

(34) Visscher, K.; Schnltzer, M. J.; Block, S. M. Single Kinesin Molecules Studied with a Molecular Force Clamp. Nature 1999, 400 (6740), 184–189.

(35) Azzam, O. Al; Trussell, C. L.; Reinemann, D. N. Measuring Force Generation within Reconstituted Microtubule Bundle Assemblies Using Optical Tweezers. Wiley 2021, No. February, 111–125.

(36) Laan, L.; Husson, J.; Munteanu, E. L.; Kerssemakers, J. W. J.; Dogterom, M. Force-Generation and Dynamic Instability of Microtubule Bundles. Proc. Natl. Acad. Sci. U. S. A. 2008, 105 (26), 8920–8925.

(37) Abraham, Z.; Hawley, E.; Hayosh, D.; Webster-Wood, V. A.; Akkus, O. Kinesin and Dynein Mechanics: Measurement Methods and Research Applications. J. Biomech. Eng. 2018, 140 (2), 1–11. 10.1115/1.4037886.

(38) Shukla, S.; Troitskaia, A.; Swarna, N.; Maity, B. K.; Tjioe, M.; Bookwalter, C. S.; Trybus, K. M.; Chemla, Y. R.; Selvin, P. R. High-Throughput Force Measurement of Individual Kinesin-1 Motors during Multi-Motor Transport. Nanoscale 2022, 12463–12475. 10.1039/d2nr01701f.

(39) Nitta, T.; Wang, Y.; Du, Z.; Morishima, K.; Hiratsuka, Y. A Printable Active Network Actuator Built from an Engineered Biomolecular Motor. Nat. Mater. 2021, 20 (8), 1149–1155.

(40) Selvaggi, L.; Pasakarnis, L.; Brunner, D.; Aegerter, C. M. Magnetic Tweezers Optimized to Exert High Forces over Extended Distances from the Magnet in Multicellular Systems. Rev. Sci. Instrum. 2018, 89 (4). 10.1063/1.5010788.

(41) Madariaga-Marcos, J.; Hormeño, S.; Pastrana, C. L.; Fisher, G. L. M.; Dillingham, M. S.; Moreno-Herrero, F. Force Determination in Lateral Magnetic Tweezers Combined with TIRF Microscopy. Nanoscale 2018, 10 (9), 4579–4590. 10.1039/c7nr07344e.

(42) Gosse, C.; Croquette, V. Magnetic Tweezers: Micromanipulation and Force Measurement at the Molecular Level. Biophys. J. 2002, 82 (6), 3314–3329. 10.1016/S0006-3495(02)75672-5.

(43) Bush, J.; Maruthamuthu, V. In Situ Determination of Exerted Forces in Magnetic Pulling Cytometry. AIP Adv. 2019, 9 (3). 10.1063/1.5084261.

(44) Wang, X.; Ho, C.; Tsatskis, Y.; Law, J.; Zhang, Z.; Zhu, M.; Dai, C.; Wang, F.; Tan, M.; Hopyan, S.; McNeill, H.; Sun, Y. NANOROBOTS: Intracellular Manipulation and Measurement with Multipole Magnetic Tweezers. *Sci*. Robot. 2019, 4 (28). 10.1126/scirobotics.aav6180.

(45) Kakugo, A.; Kabir, A. M. R.; Hosoda, N.; Shikinaka, K.; Gong, J. P. Controlled Clockwise-Counterclockwise Motion of the Ring-Shaped Microtubules Assembly. Biomacromolecules 2011, 12 (10), 3394–3399.

(46) Ito, M.; Kabir, A. M. R.; Inoue, D.; Torisawa, T.; Toyoshima, Y.; Sada, K.; Kakugo, A. Formation of Ring-Shaped Microtubule Assemblies through Active Self-Organization on Dynein. Polym. J. 2014, 46 (4), 220–225.

(47) Liu, H.; Spoerke, E. D.; Bachand, M.; Koch, S. J.; Bunker, B. C.; Bachand, G. D. Biomolecular Motor-Powered Self-Assembly of Dissipative Nanocomposite Rings. Adv. Mater. 2008, 20 (23), 4476–4481.

(48) Wada, S.; Rashedul Kabir, A. M.; Ito, M.; Inoue, D.; Sada, K.; Kakugo, A. Effect of Length and Rigidity of Microtubules on the Size of Ring-Shaped Assemblies Obtained through Active Self-Organization. Soft Matter 2015, 11 (6), 1151–1157.

(49) Akter, M.; Keya, J. J.; Kabir, A. M. R.; Rashid, M. R.; Ishii, S.; Kakugo, A. Functionalization of Tubulin: Approaches to Modify Tubulin with Biotin and DNA. Methods Mol. Biol. Clifton NJ 2022, 2430, 47—59.

(50) Keya, J. J.; Akter, M.; Kabir, A. M. R.; Rashid, M. R.; Kakugo, A. Construction of Molecular Robots from Microtubules for Programmable Swarming. Methods Mol. Biol. Clifton NJ 2022, 2430, 219—230.

(51) Case, R. B.; Pierce, D. W.; Hom-Booher, N.; Hart, C. L.; Vale, R. D. The Directional Preference of Kinesin Motors Is Specified by an Element Outside of the Motor Catalytic Domain. Cell 1997, 90 (5), 959–966.

(52) Wada, S.; Kabir, A. M. R.; Kawamura, R.; Ito, M.; Inoue, D.; Sada, K.; Kakugo, A. Controlling the Bias of Rotational Motion of Ring-Shaped Microtubule Assembly. Biomacromolecules 2015, 16 (1), 374–378.

(53) Kollmannsberger, P.; Fabry, B. High-Force Magnetic Tweezers with Force Feedback for Biological Applications. Rev. Sci. Instrum. 2007, 78 (11), 1–6.

(54) Tawada, K.; Sekimoto, K. Protein Friction Exerted by Motor Enzymes through a Weak-Binding Interaction. J. Theor. Biol. 1991, 150 (2), 193–200. 10.1016/S0022-5193(05)80331-5.

(55) Rashid, M. R.; Ganser, C.; Akter, M.; Nasrin, S. R.; Kabir, A. Md. R.; Sada, K.; Uchihashi, T.; Kakugo, A. 3D Structure of Ring-Shaped Microtubule Swarms Revealed by High-Speed Atomic Force Microscopy. Chem. Lett. 2023, 52, 100–104. 10.1246/cl.220491.

(56) Katira, P.; Agarwal, A.; Fischer, T.; Chen, H. Y.; Jiang, X.; Lahann, J.; Hess, H. Quantifying the Performance of Protein-Resisting Surfaces at Ultra-Low Protein Coverages Using Kinesin Motor Proteins as Probes. Adv. Mater. 2007, 19 (20), 3171–3176.

(57) VanDelinder, V.; Imam, Z. I.; Bachand, G. Kinesin Motor Density and Dynamics in Gliding Microtubule Motility. Sci. Rep. 2019, 9 (1), 1–9.

(58) Uçar, M. C.; Lipowsky, R. Collective Force Generation by Molecular Motors Is Determined by Strain-Induced Unbinding. Nano Lett. 2020, 20 (1), 669–676.

(59) Jamison, D. K.; Driver, J. W.; Diehl, M. R. Cooperative Responses of Multiple Kinesins to Variable and Constant Loads. J. Biol. Chem. 2012, 287 (5), 3357–3365. 10.1074/jbc.M111.296582.

(60) Furuta, K.; Furuta, A.; Toyoshima, Y. Y.; Amino, M.; Oiwa, K.; Kojima, H. Measuring Collective Transport by Defined Numbers of Processive and Nonprocessive Kinesin Motors. Proc. Natl. Acad. Sci. 2013, 110 (2), 501–506. 10.1073/pnas.1201390110.

(61) Nasrin, S. R.; Ganser, C.; Nishikawa, S.; Kabir, A. Md. R.; Sada, K.; Yamashita, T.; Ikeguchi, M.; Uchihashi, T.; Hess, H.; Kakugo, A. Deformation of Microtubules Regulates Translocation Dynamics of Kinesin. Sci. Adv. 2021, 7 (42), eabf2211. 10.1126/sciadv.abf2211.

(62) Lenz, M. Geometrical Origins of Contractility in Disordered Actomyosin Networks. Phys. Rev. X 2014, 4 (4), 041002. 10.1103/PhysRevX.4.041002.

(63) Rai, A. K.; Rai, A.; Ramaiya, A. J.; Jha, R.; Mallik, R. Molecular Adaptations Allow Dynein to Generate Large Collective Forces inside Cells. Cell 2013, 152 (1–2), 172–182. 10.1016/j.cell.2012.11.044.

(64) Head, D. A.; Briels, W.; Gompper, G. Spindles and Active Vortices in a Model of Confined Filament-Motor Mixtures. BMC Biophys. 2011, 4 (1), 18. 10.1186/2046-1682-4-18.

(65) Murata, S.; Toyota, T.; Nomura, S. ichiro M.; Nakakuki, T.; Kuzuya, A. Molecular Cybernetics: Challenges toward Cellular Chemical Artificial Intelligence. Adv. Funct. Mater. 2022, 32 (37).

(66) Kuzuya, A.; Nomura, S. I. M.; Toyota, T.; Nakakuki, T.; Murata, S. From Molecular Robotics to Molecular Cybernetics: The First Step Toward Chemical Artificial Intelligence. IEEE Trans. Mol. Biol. Multi-Scale Commun. 2023, 9 (3), 354–363.

(67) Xu, T.; Soto, F.; Gao, W.; Dong, R.; Garcia-Gradilla, V.; Magaña, E.; Zhang, X.; Wang, J. Reversible Swarming and Separation of Self-Propelled Chemically Powered Nanomotors under Acoustic Fields. J. Am. Chem. Soc. 2015, 137 (6), 2163–2166.

(68) Matsuda, K.; Kabir, A. M. R.; Akamatsu, N.; Saito, A.; Ishikawa, S.; Matsuyama, T.; Ditzer, O.; Islam, M. S.; Ohya, Y.; Sada, K.; Konagaya, A.; Kuzuya, A.; Kakugo, A. Artificial Smooth Muscle Model Composed of Hierarchically Ordered Microtubule Asters Mediated by DNA Origami Nanostructures. Nano Lett. 2019, 19 (6), 3933–3938.

(69) Wang, Y.; Nitta, T.; Hiratsuka, Y.; Morishima, K. In Situ Integrated Microrobots Driven by Artificial Muscles Built from Biomolecular Motors. *Sci*. Robot. 2022, 7 (69), 1–13.

(70) Ishii, S.; Akter, M.; Keya, J. J.; Rashid, M. R.; Afroze, F.; Nasrin, S. R.; Kakugo, A. Purification of Tubulin from Porcine Brain and Its Fluorescence Dye Modification. Methods Mol. Biol. Clifton NJ 2022, 2430, 3–16.

(71) Castoldi, M.; Popov, A. V. Purification of Brain Tubulin through Two Cycles of Polymerization-Depolymerization in a High-Molarity Buffer. Protein Expr. Purif. 2003, 32 (1), 83–88.

(72) Peloquin, J.; Komarova, Y.; Borisy, G. Conjugation of Fluorophores to Tubulin. Nat. Methods 2005, 2 (4), 299–303.

(73) Früh, S. M.; Steuerwald, D.; Simon, U.; Vogel, V. Covalent Cargo Loading to Molecular Shuttles via Copper-Free “Click Chemistry.” Biomacromolecules 2012, 13 (12), 3908–3911.

## Reference

(1) Kollmannsberger, P.; Fabry, B. High-Force Magnetic Tweezers with Force Feedback for Biological Applications. Rev. Sci. Instrum. 2007, 78 (11), 1–6.

(2) Rashid, M. R.; Ganser, C.; Akter, M.; Nasrin, S. R.; Kabir, A. M. R.; Sada, K.; Uchihashi, T.; Kakugo, A. 3D Structure of Ring-Shaped Microtubule Swarms Revealed by High-Speed Atomic Force Microscopy. Chem. Lett. 2023, 52 (2), 100–104. 10.1246/cl.220491.

(3) Katira, P.; Agarwal, A.; Fischer, T.; Chen, H.-Y.; Jiang, X.; Lahann, J.; Hess, H. Quantifying the Performance of Protein-Resisting Surfaces at Ultra-Low Protein Coverages Using Kinesin Motor Proteins as Probes. Adv. Mater. 2007, 19 (20), 3171–3176. 10.1002/adma.200701982.

